# tappAS: a comprehensive computational framework for the analysis of the functional impact of differential splicing

**DOI:** 10.1101/690743

**Authors:** Lorena de la Fuente, Ángeles Arzalluz-Luque, Manuel Tardáguila, Manuel Tardáguila, Héctor del Risco, Cristina Martí, Sonia Tarazona, Pedro Salguero, Raymond Scott, Ana Alastrue-Agudo, Pablo Bonilla, Jeremy Newman, Lauren McIntyre, Victoria Moreno-Manzano, Ana Conesa

**Author notes:** Bioinformatics Unit, IIS Fundación Jiménez Díaz, Madrid, Spain. Human Genetics Department, Wellcome Trust Sanger Institute, Hinxton (Cambridge), UK. These authors contributed equally to this work (Lorena de la Fuente, Manuel Tardáguila). These authors jointly supervised this work (Victoria Moreno-Manzano, Ana Conesa).

## Abstract

Traditionally, the functional analysis of gene expression data has used pathway and network enrichment algorithms. These methods are usually gene rather than transcript centric and hence fall short to unravel functional roles associated to posttranscriptional regulatory mechanisms such as Alternative Splicing (AS) and Alternative PolyAdenylation (APA), jointly referred here as Alternative Transcript Processing (AltTP). Moreover, short-read RNA-seq has serious limitations to resolve full-length transcripts, further complicating the study of isoform expression. Recent advances in long-read sequencing open exciting opportunities for studying isoform biology and function. However, there are no established bioinformatics methods for the functional analysis of isoform-resolved transcriptomics data to fully leverage these technological advances. Here we present a novel framework for Functional Iso-Transcriptomics analysis (FIT). This framework uses a rich isoform-level annotation database of functional domains, motifs and sites –both coding and non-coding- and introduces novel analysis methods to interrogate different aspects of the functional relevance of isoform complexity. The Functional Diversity Analysis (FDA) evaluates the variability at the inclusion/exclusion of functional domains across annotated transcripts of the same gene. Parameters can be set to evaluate if AltTP partially or fully disrupts functional elements. FDA is a measure of the potential of a multiple isoform transcriptome to have a functional impact. By combining these functional labels with expression data, the Differential Analysis Module evaluates the relative contribution of transcriptional (i.e. gene level) and post-transcriptional (i.e. transcript/protein levels) regulation on the biology of the system. Measures of isoform relevance such as Minor Isoform Filtering, Isoform Switching Events and Total Isoform Usage Change contribute to restricting analysis to biologically meaningful changes. Finally, novel methods for Differential Feature Inclusion, Co-Feature Inclusion, and the combination of UTR-lengthening with Alternative Polyadenylation analyses carefully dissects the contextual regulation of functional elements resulting from differential isoforms usage. These methods are implemented in the software tappAS, a user-friendly Java application that brings FIT to the hands of non-expert bioinformaticians supporting several model and non-model species. tappAS complements statistical analyses with powerful browsing tools and highly informative gene/transcript/CDS graphs.

We applied tappAS to the analysis of two mouse Neural Precursor Cells (NPCs) and Oligodendrocyte Precursor Cells (OPCs) whose transcriptome was defined by PacBio and quantified by Illumina. Using FDA we confirmed the high potential of AltTP regulation in our system, in which 90% of multi-isoform genes presented variation in functional features at the transcript or protein level. The Differential Analysis module revealed a high interplay between transcriptional and AltTP regulation in neural development, mainly controlled by differential expression, but where AltTP acts the main driver of important neural development biological mechanisms such as vesicle trafficking, signal transduction and RNA processing. The DFI analysis revealed that, globally, AltTP increased the availability of functional features in differentiated neural cells. DFI also showed that AltTP is a mechanism for altering gene function by changing cellular localization and binding properties of proteins, via the differential inclusion of NLS, transmembrane domains or DNA binding motifs, for example. Some of these findings were experimentally validated by others and us.

In summary, we propose a novel framework for the functional analysis of transcriptomes at isoform resolution. We anticipate the tappAS tool will be an important resource for the adoption of the Functional Iso-Transcriptomics analysis by functional genomics community.

## Introduction

One of the most exciting aspects of transcriptome biology is the contextual adaptability of eukaryotic transcriptomes and proteomes by Alternative Splicing (AS), Alternative PolyAdenylation (APA), and Alternative Transcription Start Sites (ATSS) mechanisms, jointly referred to as Alternative Transcript Processing (AltTP). These three processes determine which transcripts (aka, isoforms) are produced for a given gene. Alternate transcripts may differ in structure and in function, as well as in cell specificity, and within cell spatio-temporal deployment.

The study of AltTP has experimentally been addressed either via molecular characterization of the functionality of specific isoforms from single genes^1,2^, or by computationally approaches aiming to find global patterns and infer their potential biological significance *in silico*^3,4^. Computational AltTP analysis has focused on the study of processing *events*, namely exon spiking, intron retention, alternative transcript start (TSS) and termination sites (TTS), nonsense-mediated decay (NMD) and changes in the inclusion/exclusion levels of different exons^5–8^. In parallel, molecular studies have been conducted to understand the mechanisms behind the dynamic changes in event patterns, identifying a large number of RNA binding proteins as regulators of AltTP^9–14^. In response to the recognition of the biological importance of AltTP, bioinformatics tools have been developed to analyze the structural and regulatory aspects of AltTP events and have contributed to the description and understanding of AltTP (reviewed in^15^).

While some discrepancy exist on the actual functional role of transcript isoform diversity^16,17^. AltTP has been proven to be implicated in differentiation^18–20^, tissue identity^21,22^, development^13,23^, stress response^24^ and disease^25–28^. Beyond these well known effects, several studies have shown enrichment of spliced exons in disordered regions mediating new protein interactions^29^ and remodeling of protein-protein interaction in a tissue-specific manner^30,31^. In other work, AS was shown to regulate domains leading to the rewiring of PPI networks in cancer^32^. Similarly, APA has been postulated as a mechanism to escape microRNA regulation by shortening 3’ UTR regions^33,34^, alternative TSS are believed to regulate the inclusion of Upstream Open Reading Frames (uORFs) that control translational rates^35–37^ and NMD has been proposed to regulate gene expression in cancer and neural systems^38,39^.

Traditionally, computational approaches such as enrichment and network analysis have been used to study the functional aspects of transcriptional changes^40–43^ and these have been instrumental for the characterization of transcriptome biology. However, these methods operate at the gene level and are not adapted to study the functional readout of AltTP. Much of the work done to answer transcriptome-wide questions on the functional role of AltTP has involved *ad hoc* computational pipelines applied to specific biological systems or address only particular types of events^44–49^. Recently, Exon Ontology^50^ was proposed as a resource to study functional enrichment of exon sets based on their annotation with protein functional domains. Using this tool, authors were able to show different molecular functionalities directly associated to changes in exon inclusion levels between epithelial and mesenchymal cells. However, this analysis does not reveal how transcripts combine exons to provide distinct functional elements, nor addresses the analysis of regulatory signals at alternative UTRs. In general, the field lacks computational tools tailored to the study of the functional aspects of isoform expression regulation, limiting advances in our understanding of the functional impact of AltTP.

One important reason behind the lack of functional perspective in splicing-dedicated bioinformatics tools is the inability of RNAseq to correctly capture isoform expression^51^. Recently, third generation sequencing technologies have demonstrated their power in detecting full-length transcript^52–56^ and identifying expressed isoforms. Options for quantification are found in the combination with short-reads^55^ or the utilization of the newest high throughput instruments. As more scientists engage in expression studies that use these new platforms with the goal of identifying differences between conditions in isoform usage, there is a growing need of tools to easily and quickly interpret isoform differences in the context of their potential functional impact.

Here we present a novel computational framework for the study AltTP from a functional perspective, introducing the Functional Iso-Transcriptomics (FIT) analysis approach. This framework uses a rich isoform-level annotation database of functional domains, motifs and sites –both coding and non-coding-, that are mined by novel analysis methods that interrogate different aspects of the functional load associated to isoform complexity and expression regulation. These methods are implemented in the software tappAS (http://tappas.org), a user-friendly Java application that brings FIT to the hands of transcriptome scientists by supporting several model and non-model species. tappAS complements statistical analyses with powerful browsing tools and highly informative gene/transcript/CDS graphs. As a proof of principle, we applied tappAS to the analysis of two mouse neural cell types, Neural Precursor Cells (NPCs) and Oligodendrocyte Precursor Cells (OPCs), whose transcriptome was defined by PacBio and quantified by Illumina^57^. tappAS easily recapitulates a great deal of the existing knowledge on AltTP function, as well as provide new functional insights. We anticipate that the tappAS framework will be widely applied in a variety of fields, and that its user-friendliness will promote the adoption of the FIT approach by researchers with different levels of computational skills.

## Results

### tappAS is a comprehensive tool to investigate potential functional consequences of AltTP

tappAS analyses use a species-specific gff3-like file containing isoform-level, positionally-resolved, annotation features (see Methods). These labels describe functional motifs, domains and sites both at the CDS and the UTRs of transcripts, and are generated via the integration species-available databases and sequence-based prediction tools that gather functional and structural data. For our mouse example, 20 functional categories were retrieved (Supplementary Table 1). tappAS joins transcript-level expression data with this extensive annotation database and a wide array of traditional and novel analysis algorithms (Table 1) to create a comprehensive framework for the study of the functional impact of AltTP.

**Table 1:**
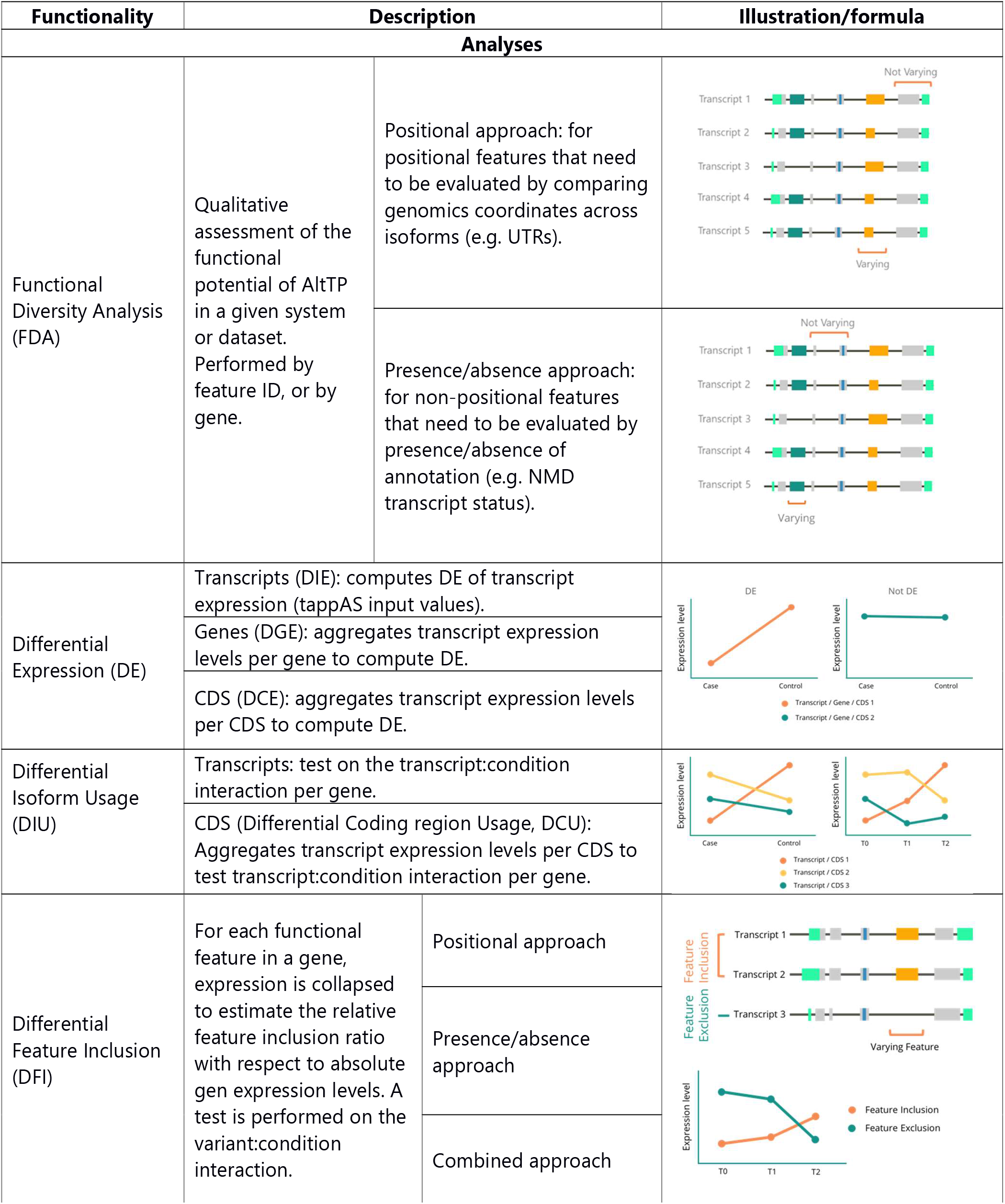
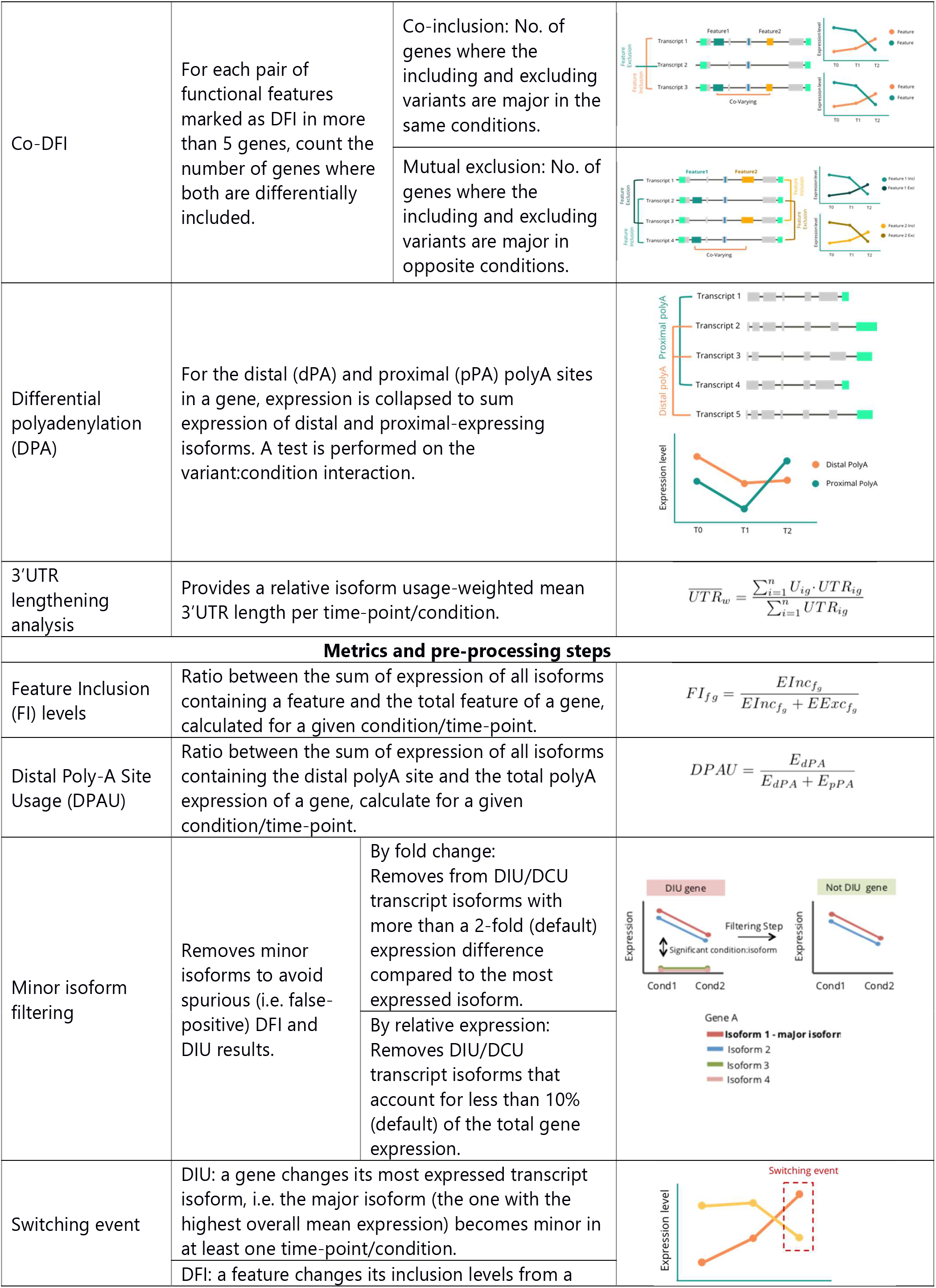
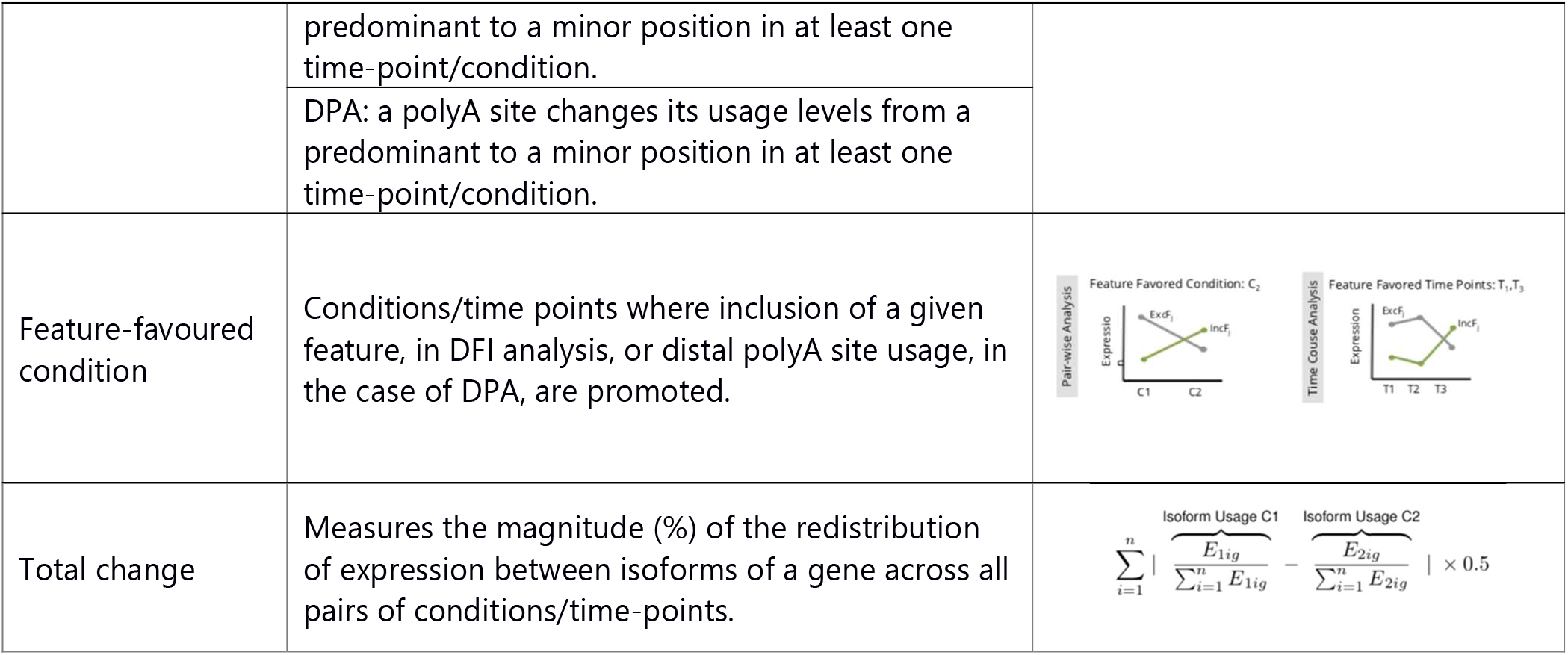
Main analyses and metrics of tappAS.

tappAS analysis can be divided into three Modules, each one targeting a different aspect in the study of AltTP biology (Figure 1). Module I includes Functional Diversity Analysis (FDA), which evaluates the functional regulatory potential of AltTP by interrogating the varying status of individual features across isoforms of the same gene (Figure 1). This includes analysis by gene (assessing the varying status of individual genes for each functional feature category) and by feature ID (assessing the number of genes for which a particular feature is differentially present across isoforms). Depending on the feature, varying status is evaluated by genomic position (positional approach) or by presence/absence (presence approach) (Table 1, Methods). Module II can be used to understand the relative contribution of transcriptional and post-transcriptional regulation in the system under study by comparing Differential Isoform Usage (DIU, transcript level) or Differential Coding sequence Usage (DCU, protein level) with Differential Gene Expression (DGE) results, and by performing subsequent enrichment analyses. Finally, Module III includes methods to assess the context-dependent differential inclusion of annotated functional elements: Differential Feature Inclusion (DFI) of coding and non-coding elements, Differential PolyAdenylation (DPA) and 3’UTR lengthening analysis (3UL). Furthermore, a subsequent co-Differential Feature Inclusion (co-DFI) analysis can detect sets of features that are coordinately included. DPA and 3’UTR lengthening analyses can be combined to study which genes are regulated via alternative polyadenylation (APA) and 3’UTR length. Importantly, any of the tappAS outputs described above can be coupled to Functional Enrichment^58^ and Gene-Set Enrichment^59^ analyses based on any of the functional categories included in tappAS annotation. Finally, tappAS’ displays all annotated features as gene, transcript and protein graphical maps enabling for a visual evaluation of isoforms and their functional components. For more details on the methodology behind these analyses, see Online Methods.

**Figure 1:**
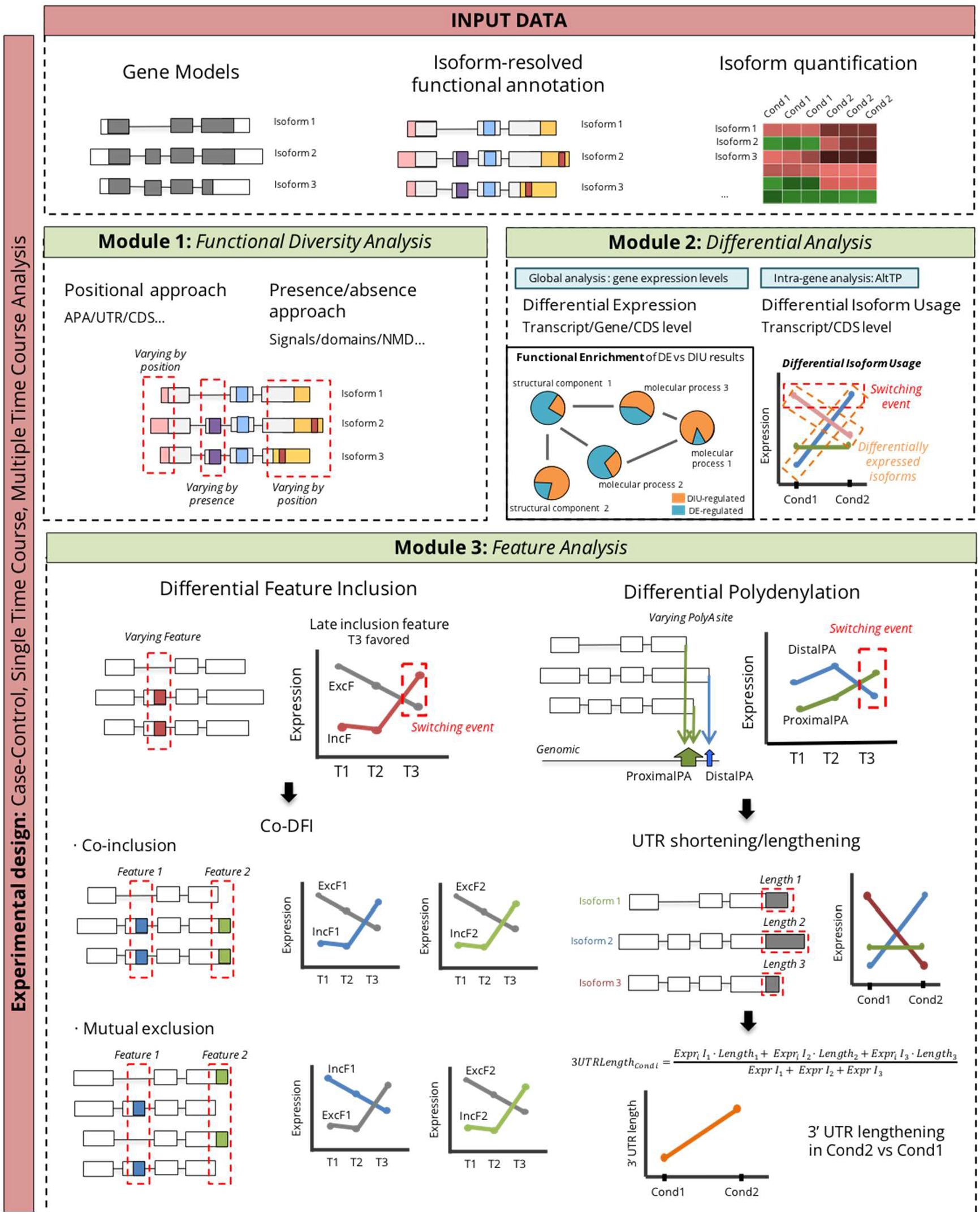
Overview of tappAS modules for Functional Iso-Transcriptomics Analysis. Module 1 contains a novel qualitative approach to evaluate functional diversity of alternative isoforms. Module 2 implements Differential Expression and Differential Isoform Usage analyses to discriminate AltTP (post-transcriptional) from transcriptional regulation mechanisms. Module 3 includes newly-developed approaches to measure the functional impact of AltTP as changes in the inclusion of functional features, polyA site usage and UTR length.

### Functional Diversity Analysis

One fundamental question about AltTP is how post-transcriptional regulation imprints functional complexity to transcriptomes. The potential of AltTP mechanisms to regulate gene function largely depends on whether transcript isoforms contain variation in their functional elements. In this case, modifications in their expression levels can effectively modulate functional changes. Applied to our murine neural transcriptomes, tappAS FD analysis identified ~70% of 2,341 multi-isoform genes that varied in the predicted proteins (Figure 2A, CDS variability). Variability at 3’ and 5’ UTR lengths occurred in ~ 60% of the genes (Figure 2A). The vast majority (78%) of UTR-varying genes also had CDS variation, suggesting that protein diversity may be coupled to RNA regulatory diversity. To illustrate, Figure 2C shows an example of a gene detected by tappAS as Alternative PolyAdenylation (APA), 5’UTR, 3’UTR and CDS-varying.

**Figure 2:**
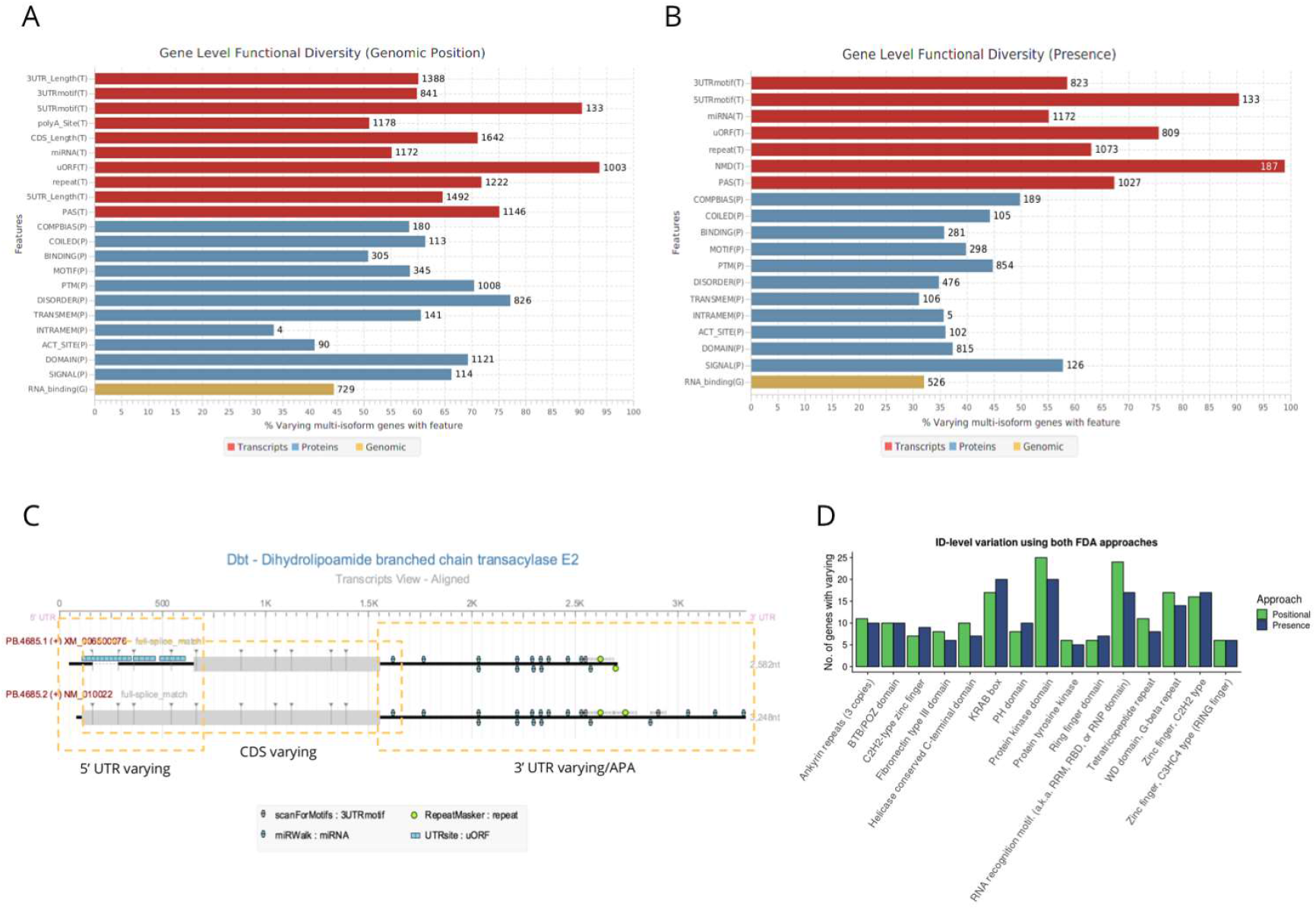
Functional Diversity Analysis (FDA) results. A) FDA results summary using the positional approach. The % of multi-isoform genes with the annotated feature in which at least one isoform is varying is shown. The numbers above the bars indicate the total no. of varying genes for that category. B) FDA results summary using the presence/absence approach. C) tappAS graphical representation of the transcript-level annotation for the *Dbt* gene, where 5’UTR, CDS and 3’UTR/Alternative polyadenylation variation can be observed. D) Comparison of position vs presence/absence approach FDA results for the ID-level analysis of variation in PFAM domains. Top-15 domain families ranked by total number of varying genes shown.

Nonsense-mediated decay (NMD) had the highest variation rate among transcript-level features, 95% (Figure 2B). Moreover, nearly all genes with NMD transcripts expressed protein-coding counterparts, indicating that NMD-targeted isoforms are co-expressed with functional isoforms in our neural system, likely regulating their abundance^38,39,60^. UTR-motif annotated genes showed a presence/absence varying rate of 55% and 90% for 3’ and 5’ UTR motifs, respectively (Figure 2B), and GU-rich elements (GREs) were the most significantly varying 3’UTR motif types (Supplementary Table 2). GREs have been associated to the stabilization of mRNAs^61^ and also have been reported as targets of RNA-binding proteins (RBPs) such as CELFs^62^. Among the set of 160 genes with differential inclusion of GRE elements in our neural system, tappAS identified splicing regulators such as *Rbm4* (Supplementary Figure 1A), involved in neurogenesis of the mouse embryonic brain^63^, and *Tcf12* (Supplementary Figure 1B), known to play an important role in the control of proliferating neural stem cells and progenitor cells during neurogenesis^64^.

FDA also identified la large number of miRNAs. An enrichment test was used to rank miRNAs that we more frequently varying at 3’UTRs (Supplementary Table 3). Interestingly, the top-five most significantly varying miRNAs include miR-335-3p, known to associate with oligodendrocyte differentiation^65^, and mir-590-3p, which responds to retinoic acid and is strongly associated to proliferation and differentiation processes^66^. Since our NPCs and OPCs constitute differentiating primary cells, these results point towards a potential isoform-specific layer of expression regulation in neural differentiation via gain and loss of miRNA binding sites due to AltTP.

Regarding presence/absence FD analysis of protein-level features, signal peptides have the highest varying rate, followed by compositional bias regions and post-translational modifications (PTMs) (Figure 2B). However, most features involving functional variability within coding sequences are best studied via the FD positional approach (Table 1, Figure 2A), which reports cases where a functional feature is partially disrupted, suggesting functional modulation changes. Hence, considering positional variation, Intrinsically Disordered regions (IDRs) and PFAM domains present the highest rates of differential inclusion in multi-isoform genes annotated for these feature categories (~78% and ~70%, respectively; Figure 2A) when compared to presence/absence variation (Figure 2B). IDRs have been reported to be frequently present in transcript regions affected by AltTP^29,67,68^.

To understand which PFAM domain types have higher positional than presence varying rates, we interrogated this category at the ID level. Figure 2D shows the top-15 PFAM domains ranked by varying rate in our data, using both the positional and presence approaches. We observe that zinc fingers and KRAB-box domains tend to be totally contained in AS exons, as varying rates using the presence and positional approaches are only slightly different. Hence, domain skipping in these cases will result in elimination from the protein, while Kinase and RNA binding domains stand out at the positional FD analysis, indicating that AltTP mechanisms tend to partially disrupt these domains, possibly causing partial loss/change of function.

In summary, tappAS’ FD analysis successfully catalogues the transcriptome’s potential for AltTP-mediated functional diversity and, in our mouse neural system, reveals that ~90% of multi-isoform genes have protein or transcript-level functional features that vary across isoforms.

### Multi-layered Analysis of Alternative Transcript Processing

Transcriptional and post-transcriptional (AltTP) regulation are regulatory mechanisms that either control total expression levels or differences in the relative isoform proportions, both contributing to regulate gene function. tappAS’ Differential Module (Figure 1) is designed to dissect and compare these two regulatory layers.

Figure 3A shows the intersection of differential analysis results for our neural dataset. tappAS identified 1,205 genes differentially expressed between NPCs and OPCs (FDR<0.05, FC>1.5), while only 291 of them were also regulated by AltTP mechanisms, as revealed by DIU analysis (FDR<0.05). Interestingly, although these results showed that most DE multi-isoform genes were regulated exclusively at the transcriptional level, a group of 247 genes were solely affected by AltTP, meaning that ~50% of DIU genes underwent a redistribution of expression among their isoforms with no significant change in gene-level expression (example in Figure 3B). This suggests independent AltTP and gene expression regulatory mechanisms operating in our neural system. However, when a filter on isoform low relative abundance was applied (<10% of total gene expression), 110 genes lost DIU status, revealing that a fraction of DIU calls is composed by transcripts that barely contribute to total gene expression, and might not be functionally relevant (Supplementary Figure 2A). After DCU analysis (Table 1, FDR<0.05), we identified a group of 135 genes where differential usage of isoforms did not involve changes in coding sequence usage (see examples in Supplementary Figure 2B, comprehensive results in Supplementary Table 4). Finally, among 279 genes detected by both DIU and DCU analyses after filtering (and therefore significantly affected by AltTP), a relevant 35% undergo a major isoform switch (see Methods), (Table 1) between NPCs and OPCs (Figure 3A), meaning that a significant fraction of isoform usage differences between both cell types have the potential for a strong functional impact. In order to identify the most significantly AltTP-regulated candidates, we used joint evaluation of total usage change (Table 1, Online Methods), which constitutes a quantitative measure of DIU (i.e. the degree of isoform usage change for a given gene across conditions), together with the identification of isoform switching events (Figure 3C). Specifically, most genes with isoform switching have total change >20%. Hence, switching can be used as criteria to prioritize candidates where AltTP has potentially higher impact on the functionality of the gene and are more interesting for experimental validation.

**Figure 3:**
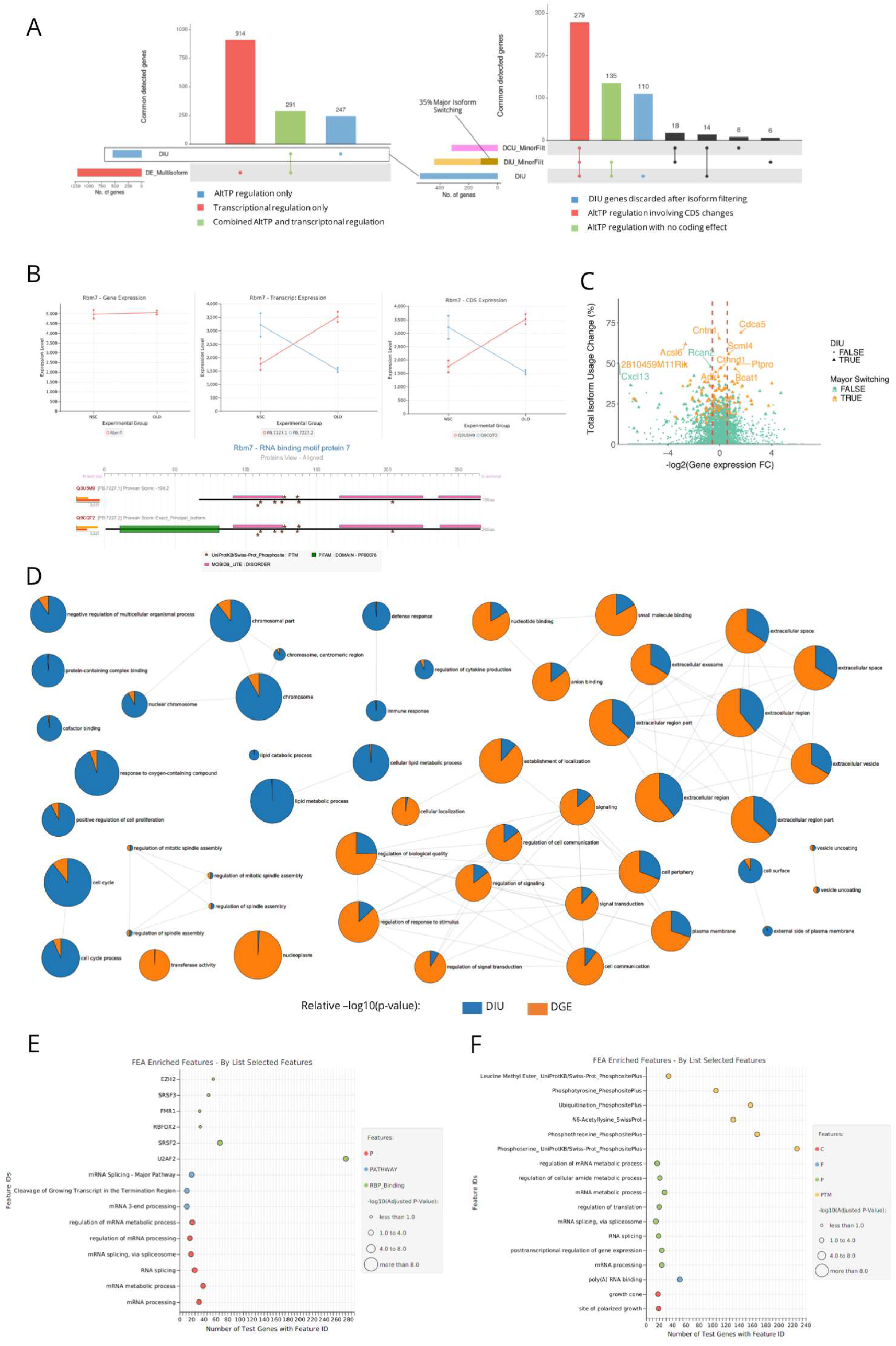
Combined analysis of differential gene expression and AltTP in tappAS. A) UpSet plot showing intersections of DE vs DIU (left) and DIU vs DCU results, with and without minor isoform filtering. Horizontal bars correspond to the total set of genes detected as significantly DE or DIU. Matrix points indicate the evaluated intersection, and vertical bars indicate their size. Legends detail the biological importance of each intersecting set of genes. B) From left to right, gene, transcript and protein-level expression charts for the *Rbm7* gene in our system, and tappAS graphical representation of its protein-level annotation. While there are no changes in gene expression level (not DE), the gene presents both differential isoform and coding sequence usage. C) Total usage change (i.e. expression redistribution between isoforms) vs log-transformed values of gene expression fold change between cell types. Genes with a major isoform switch are represented in orange. Labels are assigned to genes with the highest total usage change, indicating also whether they undergo major isoform switching. D) Multi-Dimensional Gene Set Enrichment Analysis of genes ranked by DE and DIU p.value. Nodes correspond to GO-terms obtained by selecting the top-25 terms ranked by significance in the DE enrichment and the top-25 terms ranked by significance in the DIU enrichment. Pie chart area represents DE and DIU regulation, and corresponds to relative −log10(p-value). E) and F) Functional Enrichment of DIU (E) and DCU (F) genes (Fisher’s Exact Test, with Benjamini-Hochberg multiple testing correction, minor isoform filtering: proportion < 10%) using DE genes as background. Dot color indicates the functional category of the feature, while dot size indicates significance.

To evaluate the potential functional impact of AltTP relative to gene expression regulation, tappAS includes Functional Enrichment algorithms operating on all available functional databases and sets of differential features. For example, Gene Ontology-based Multi-Dimensional Gene Set Enrichment Analysis^69^ of genes ranked by DE and DIU p-value is effective to directly compare enriched functions controlled by either mechanism. Figure 3D shows the top 25 enriched GO terms in this analysis. In this tappAS representation we readily appreciate that transcriptional regulation dominates in some important functions required for differentiation, as shown by preferential enrichment in cell cycle, spindle and chromosome-related terms. DE-regulation is also the main driver of some processes related to oligodendrocyte function, such lipid metabolism, likely related to myelination (Figure 3D). On the contrary, preferential regulation by AltTP is present for core of terms related to vesicle transport, in line with the known role of vesicle trafficking for polarity establishment and myelination^70–72^, and with previous reports of splicing regulation of vesicle transport^73^, also during differentiation processes^74^. A second group of terms related to signalling and cell communication also appears highly regulated by DIU, which together suggest high importance of AltTP in the response to extracellular signals, as recently reported^75^. This is particularly relevant to our system given that external stimuli are known to be involved in development^76^ and require activation of signaling pathways for an integrated differentiation response. In addition, the strong DIU regulation of terms such as *neuron projection* (Supplementary Figure 3), *plasma membrane* and *cell periphery* (Figure 3D) is in agreement with the established role of cell polarity and shape for NSC differentiation towards oligodendrocytes and the successful establishment of the myelin sheath^70^. Moreover, analysis of neural specific terms revealed that, while the regulatory terms (involving neuron survival and neurogenesis regulation) are mainly DE-regulated, the underlying differentiation processes (neurogenesis and neuron differentiation-related processes) predominantly involve DIU genes (Supplementary Figure 3). This suggests a transcriptional control of differentiation regulators that trigger differentiation processes that are in turn mostly AltTP-mediated. These results therefore point towards a strong interplay between gene expression and AltTP regulation, where the synergies between both, as well as each individual effects, are the ultimate drivers of biological processes that are key to neural development.

Finally, to deepen into the cellular functionalities solely regulated by AltTP, we used tappAS to calculate enrichment of DIU and DCU genes using the set of DE genes as background. As well as targets of several RNA binding proteins, we found significant enrichment of processes involved in 3’-end mRNA processing, RNA binding and mRNA splicing (Figure 3D), pointing towards a high degree of self-regulation of the post-transcriptional machinery in our system. Indeed, genes from several splicing regulator families, such as Ser/Arg-rich splicing factors (*Srsf5, Srsf10*), Muscleblind-like proteins (*Mbnl1, Mbnl2*) and RNA-binding motif proteins (*Rbm5, Rmb7*) undergo significant differential isoform/protein usage in our system (Supplementary Figure 4, Supplementary Figure 6A). Additionally, the analysis indicated enrichment of DCU genes for cellular components and processes associated to neural development, such as neurite/axon outgrowth (*growth cone*, FDR=0.002; *site of polarized growth*, FDR=0.001)(Figure 3E), showing that the analysis of protein isoform changes may reveal interesting processes that remain hidden when solely looking at transcript usage. Moreover, significant enrichment was found for NLSs (FDR=0.02), indicating that differential coding sequence usage may change the subcellular localization of the resulting protein, and several PTMs (Phosphoserine, FDR=0.02; Phosphothreonine, FDR=0.02; Acetyl-Lysine, FDR=0.05), suggesting that AltTP may be related to post-translational modulation of protein function.

In conclusion, combining tappAS Differential and Enrichment modules allows disentangling the contribution of transcriptional and post-transcriptional regulation to transcriptome changes. In our proof of concept experimental system, both mechanisms affect to shared and specific processes jointly shaping the cell type differences.

### Feature-level Analysis of AltTP and Differential Isoform Usage

To investigate how functional features are included/excluded due to differential usage of isoforms and AltTP, we applied tappAS’ Differential Feature Inclusion (DFI) analysis. Differentially included features between NPC and OPC were identified in 526 genes, including ~83% of previously detected DIU genes, indicating that our framework recapitulates post-transcriptional regulation with changes in the functional properties of transcripts and proteins. Features positive for DFI were found distributed along all considered categories (Figure 4A), although a significant relative enrichment was found for uORFs (Fisher’s exact test (FET) p-value=5.25e-121), RNA binding protein (RBP) binding sites (FET p-value=2.46e-07), compositional bias regions (FET p-value=4.06e-03) and IDRs (FET p-value 5.02e-03). Gene level DFI also indicated IDRs and 5’UTR elements (particularly uORFs) as significantly differentially included (Figure 4B).

**Figure 4:**
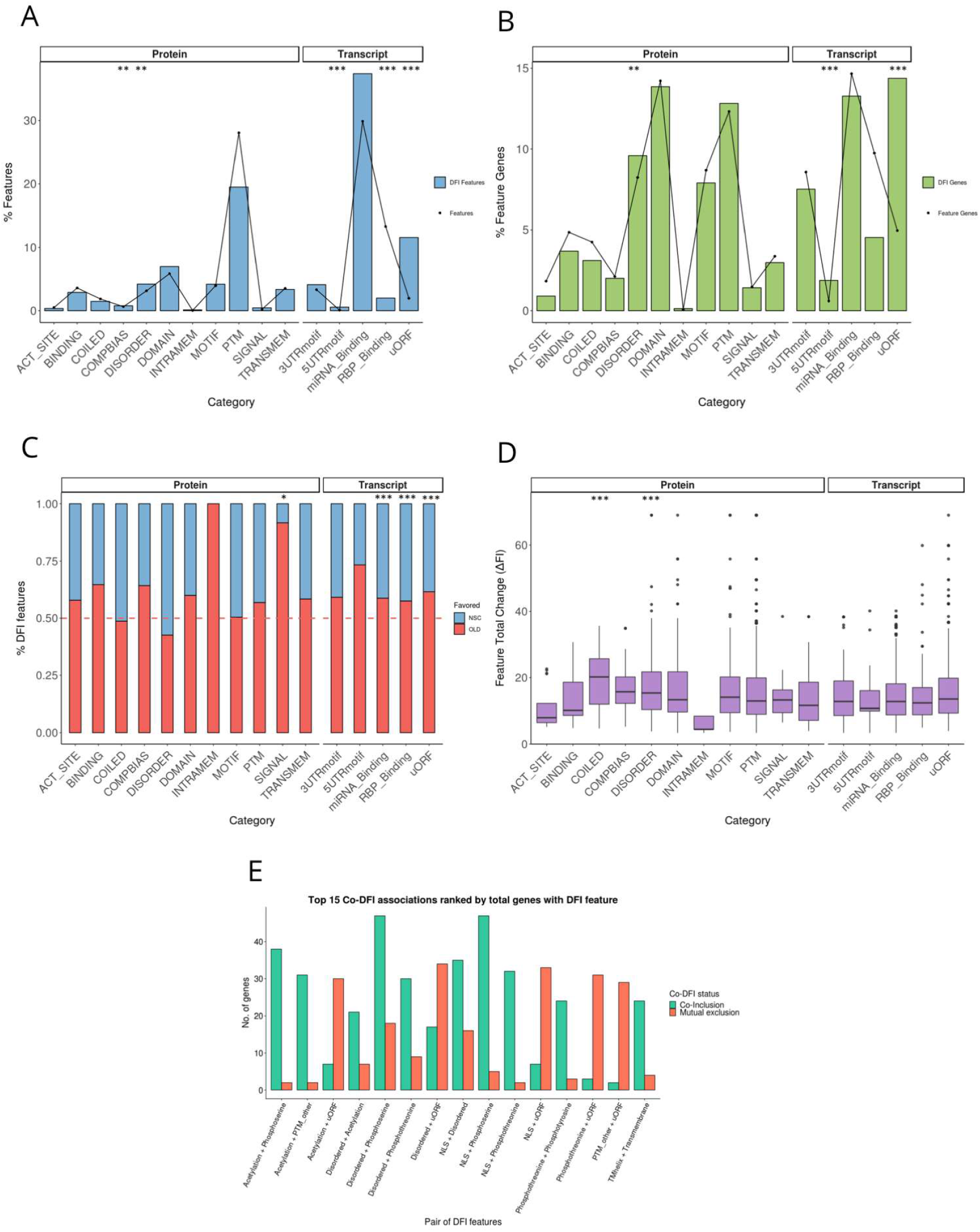
Summary of DFI results. A) Distribution (%) of features annotated in the transcriptome (dots) vs differentially included features revealed by the analysis (bars). The relative over-representation of DFI features in specific categories is evaluated by Fisher Exact tests and corrected for multiple testing using the Benjamini-Hochberg method. Significant categories are marked by asterisks (*). B) Distribution (%) of genes annotated for each feature category (dots) vs genes with differentially included features (bars). Significance (*): FET with Benjamini-Hochberg correction. C) Distribution (%) of differentially included features according to the cell type in which the inclusion of the feature is favored. Categories enriched in cell type specific inclusion were captured using a Binomial test with probability = 0.5 and Benjamini-Hochberg multiple-testing correction. D) Differences in inclusion levels for each feature category across cell types measured by feature total change (ΔFI). Differential distribution across categories tested with the non-parametric Kruskal test. Significance scale: (***) p < 0.001; (**) p < 0.01; (*) p < 0.05. E) Top 15 co-DFI associations ranked by total genes with both features marked as DFI. Bar color indicates the number of genes where features are co-included in the same conditions (co-inclusion) or in opposite conditions/groups (mutual exclusion). F) Summary of features found to be significantly DFI using different comparison strategies.

Moreover, we found feature gain to be more frequent in OPCs when compared to NPCs (Figure 4C), which can be interpreted as AltTP promoting the incorporation of functional properties as cells differentiate. For example, we observed OPC-specific inclusion of signal peptides (Binomial test, probability of success = 0.5, BiTest FDR = 2.10e-02), as well as of miRNA binding sites (BiTest FDR = 3.85e-08), uORFs (BiTest FDR = 5.85e-31) and RBP binding sites (BiTest FDR = 4.09e-04), which may indicate a 3’ UTR lengthening trend in OPCs vs NPCs. Remarkably, when comparing (absolute) differences in feature inclusion rates between the two cell types, we found them to amount no more than 20% for most categories, suggesting that in our system AltTP acts as mechanism for the functional fine-tuning of gene products. Nevertheless, we found significant differences in feature total change (ΔFI, see Methods) across functional categories, being coiled regions and IDRs the protein domains with the highest change in inclusion levels between cell types (Figure 4D, Mann-Whitney test, disordered FDR = 3.63e-07, coiled FDR= 5.43e-06).

In total, 526 genes were significant for DFI analysis, and many of these differentially included features related to binding properties and cellular localization. For example, we found a significant number of genes with isoforms differentially including Nuclear Localization Signals (n = 89), possibly regulating their switch between nucleus and cytosol as cell differentiate. This is the case of the *Ctnnd1* gene encoding p120, a well-known component of the β-catenin signaling pathway, an important process in the differentiation of NPCs to OPCs^77,78^. tappAS predicted that *Ctnnd1* possesses an NLS motif in two of its alternative transcripts that appears due to exclusion of exon 10 (Figure 5A). We found *Ctnnd1* NLS-containing isoforms to be strongly downregulated in NPCs, while an isoform switching event leads to a significant increase in their expression levels in OPCs. Western blot analysis of *Ctnnd1* confirmed a localization change in OPCs and the increase of nuclear levels of the protein, while a cytoplasmic retention was observed in NPCs (Figure 5D). Similarly, tappAS found differential expression for the NLS of RBP *Mbnl1*, an important neural splicing factor. tappAS analysis indicates that nuclear MBNL1 isoforms are significantly favored in NPCs with respect to OPCs (DIU p-value = 0.0018; Supplementary Figures 5A-C), and Western blot analyses confirmed these observations (Supplementary Figure 5D). Finally, tappAS also detected examples of DFI affecting binding properties. Isoforms of DNA-binding protein *Mbd1* showed differential inclusion of a non-constitutive zinc finger domain (Supplementary Figure 6A), favored as differentiation progresses (Supplementary Figure 6B), and further examination suggests a potential dual mechanism that involves both differential inclusion of exon 11 in OPCs and global upregulation of *Mbd1* gene expression (Supplementary Figure 6C). In agreement, post-transcriptional processing of *Mbd1* regulating the inclusion of exon 11 zinc finger domain was recently found to be an important determinant of cell lineage in NPCs^79^.

**Figure 5:**
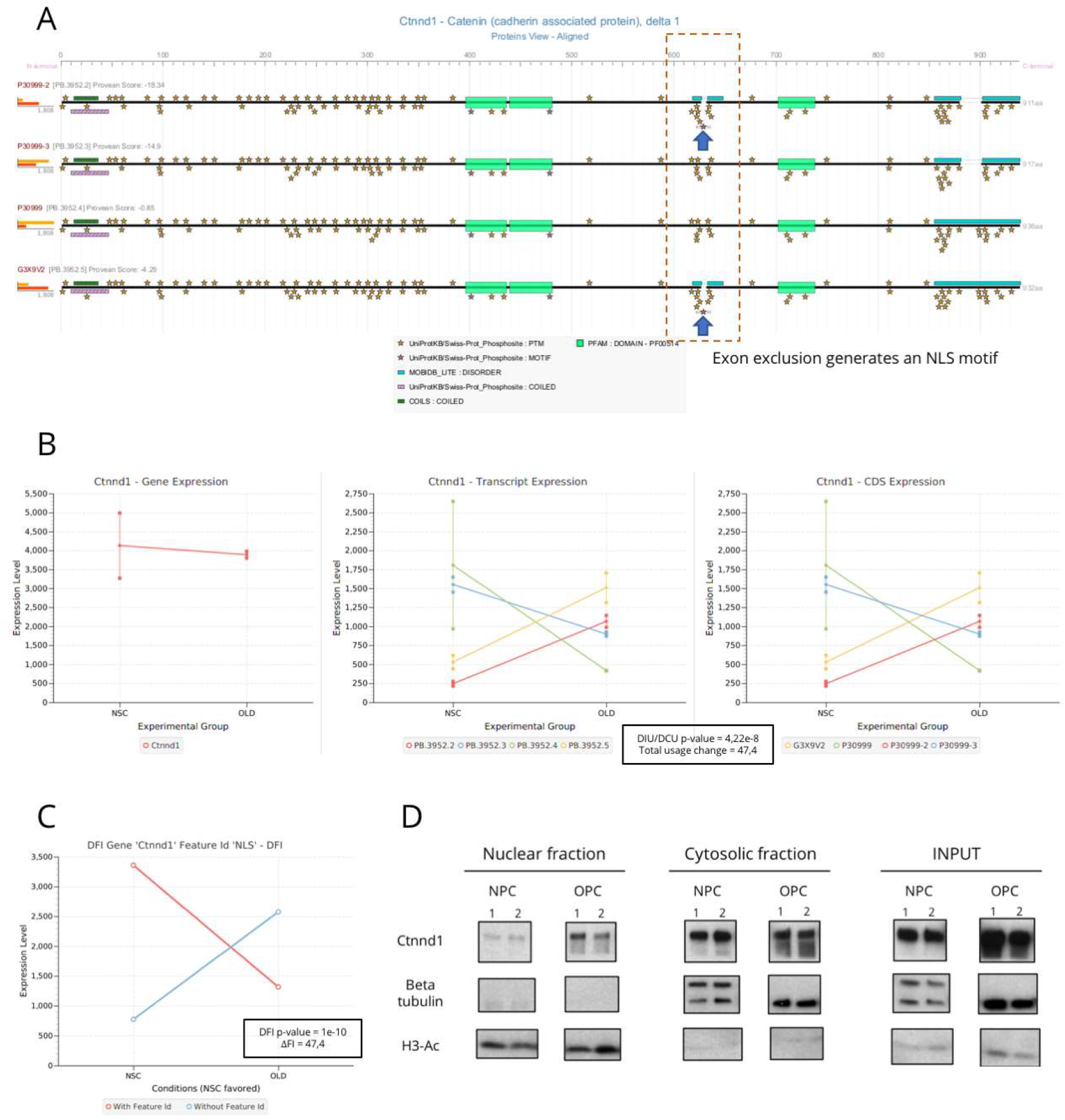
tappAS analysis results and experimental validation AltTP processing of *Ctnnd1*. A) Protein-level visualization of tappAS functional annotation for *Ctnnd1*. Exclusion of an exon causes an NLS motif to appear in the sequence. B) Gene, transcript and CDS-level expression of *Ctnnd1*. The gene is significant for both DIU and DCU, with major isoform switching of the nuclear isoforms (yellow and red) in OPCs. C) DFI analysis results for the NLS motif in *Ctnnd1*. NLS inclusion is favored in OPCs. D) Western blot analysis of *Ctnnd1* in the nuclear and cytosolic fractions of NPCs and OPCs. An increase of the nuclear expression of the protein is observed in OPCs due to differential inclusion of the NLS, while cytosolic expression remains constant.

Another interesting functionality of tappAS is the ability to investigate the coordinated inclusion of functional features, by co-DFI analysis (Figure 4E). Results revealed associations between NLS and phosphoserine residues (examples in Supplementary Figure 7A and 7B) and C2H2-type zinc finger domains (examples in Supplementary Figures 7C and 7D). Indeed, post-translational masking of NLS is a known mechanism to prevent nuclear import^80,81^. Interestingly, IDRs are also strongly co-included with phosphoserine residues, confirming their described role in the allocation of PTMs, as well as their clear association to alternatively-spliced regions (examples in Supplementary Figure 7A and 7B).

### Differential Polyadenylation

Alternative polyadenylation and differences at UTR lengths are involved in the regulation of mRNA stability, sub-cellular location, RNA protein binding and translation efficiency^82,83^. To assess the contribution of AltTP to these processes, tappAS implements Differential PolyAdenylation (DPA) and 3’UTR Lengthening (3UL) analyses (Table 1).

Applied to our experimental system, tappAS found that 17% of genes with polyA site variation across isoforms were positive for DPA (134 out of 1527, FDR < 0.001), among which ~31% (32 genes) switched their major polyA site between cell types (Figure 6A). A 56% of genes favored distal polyA site usage (DPAU) in OPCs and a significant trend towards 3’ UTR lengthening was present for OPCs (Figure 6B, Wilcoxon signed rank test, p-value = 2.267e-05). These results are consistent with our enrichment analysis (Figure 4C). Moreover, 51 genes undergoing APA regulation also had differential inclusion of miRNA binding motifs, with ~64% of DFI miRNA sites being included in OPCs. tappAS functional annotation indicated that an important number of these genes were involved in RNA processes, including *Papola, Tardbp* and *Tdrd3*, a transcriptional activator in the nucleus that is also involved in the formation of stress granules and the regulation of mRNA translation in the cytoplasm^84^. *Tdrd3* undergoes Coding Region APA, resulting in OPC upregulated forms with simultaneous inclusion of miRNAs binding sites and AU-Rich elements (ARE) at the 3’UTR (Figure 6C) and disruption of a phosphotyrosine site and an exon-junction (EJC) interacting region (Figure 6C) at the coding region. This pattern of functional regulation poses new hypothesis for the *Tdrd3* regulation by AltTP.

**Figure 6:**
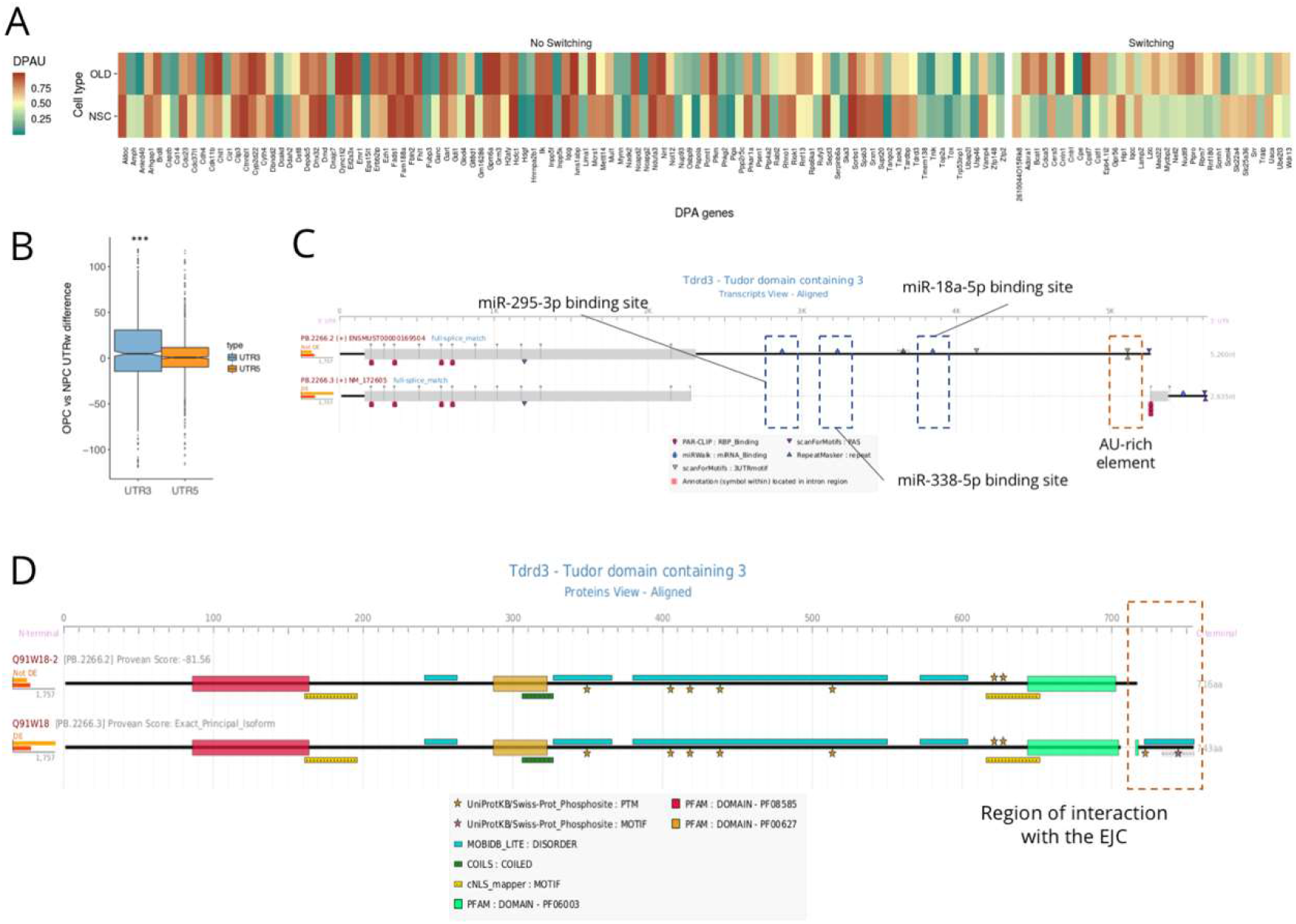
DPA results. A) Heatmap displaying DPAU levels associated to genes that are significantly DPA (FDR < 0.05) for each cell type. B) Boxplots showing the distribution of the difference in expression-weighted 3’ and 5’ UTR lengths (UTRw) in OPCs vs NPCs. C) tappAS visualization of transcript-level annotation for the *Tdrd3* gene, where Codign Region -APA induced inclusion of several miRNA binding sites as well as an AU-rich element can be observed. D) *Tdrd3* protein-level annotation, EJC binding motif variation region is squared.

## Discussion

In this work we present a novel analysis framework, implemented in the tappAS software, for the comprehensive functional analysis of isoform-resolved transcriptomes, referred here as Functional Iso-Transcriptomics (FIT). tappAS includes approaches for the analysis of the variability in functional sites at genes with multiple expressed transcripts, as well as methods to evaluate the functional impact of the context-dependent expression of alternative isoforms, and in particular to dissect which functional elements change as a consequence of differential isoform usage. We combine new analytical concepts such as FDA, DFI and U3L with more established enrichment methods to create a powerful analytical framework. This is a timely development at a moment when long-read technologies are becoming increasingly accessible, providing more accurate measurements of full-length transcripts and hence of isoform expression. However, we should highlight that tappAS is agnostic to the source of transcript models and therefore can also leverage other recently proposed strategies to improve accuracy at transcript calls such as the combination of ChIP-seq and RNA-seq data^85^ and the pre-filtering of reference isoforms based on Event Analysis^86^. Given the pace of technology, we expect that full transcript resolution and quantification will be possible in the near future. While many methods to statistically evaluate isoform expression differences do exist^5–8^, a tool specifically tailored to extract the functional readout of these isoform differences was missing, and hence tappAS comes to fill an important bioinformatics gap for the study of AltTP biology.

tappAS is designed to be a flexible framework for functional analysis of isoforms, that uses an annotation file and many options for data analysis. At present tappAS includes pre-computed gff3 files for human, mouse, fly, arabidopsis and maize. In this work we illustrate the tool with the characterization of isoform differences between two mouse neural cell types. We show that tappAS recapitulates much of the existing knowledge about this neural system, as well as of functional aspects of splicing and UTR regulation. Moreover, we show that the tappAS framework is able to propose novel functional hypothesis that can be experimentally validated, such as the alternative inclusion of NLS in proteins regulated by splicing. However, the illustrating analysis, although comprehensive, does not cover all the tappAS potentiality. Options for specifying specific sets of genes or combining multiple functional layers are available, creating endless possibilities to interrogate the data. Video tutorials at the tappAS web site (tappas.org) showcase additional functionalities of this tool. Also, as gff3 can be directly uploaded by the user, additional data could be incorporate to allow for new questions. For example, at present, no Protein-Protein interaction data or conservation scores are included in the tappAS files. Users with confident annotations at these layers can update gff3 files and easily use the tappAS framework to pose questions regarding their association with isoforms and interactions with other functional layers. Similarly, as the tool is not limited by organism, but only by the current availability of annotation, other species not yet supported in the application will benefit from tappAS as functional information becomes available.

## Acknowledgements

This work has been funded by Spanish Ministry of Education grant FPU2013/02348, Spanish MINECO BIO2015-1658-R and University of Florida Start up funds. We thank Dr. Manuel Tardáguila for assisting experimental validations and for valuable discussions on interpretation of results.

## Author contributions

LdF: Major developer of analysis methods, analyzed data, interpreted results, contributed to software implementation and testing.

AA: Created manuscript draft and validated tappAS application.

MT: Developed analysis methods, performed validation experiments and contributed to interpretation.

HdR: Major software engineer of tappAS application.

MCM: Carried out experimental procedures and validation experiments.

ST: Contributed to statistical methods development.

PS: Contributed to software implementations.

RS: Contributed to validation experiments.

AAV: Performed cell culture experiments.

PB: Contributed to validation experiments.

JRBN: Contributed to drosophila isoform annotations.

LMM: Contributed to statistical methods development and manuscript writing.

VMM: Supervised experimental work, contributed to validation of results and data interpretation.

AC. Conceived the study, supervised experimental and analytical approaches and completed manuscript draft.

## Supplementary material

**Supplementary Figure 1:**
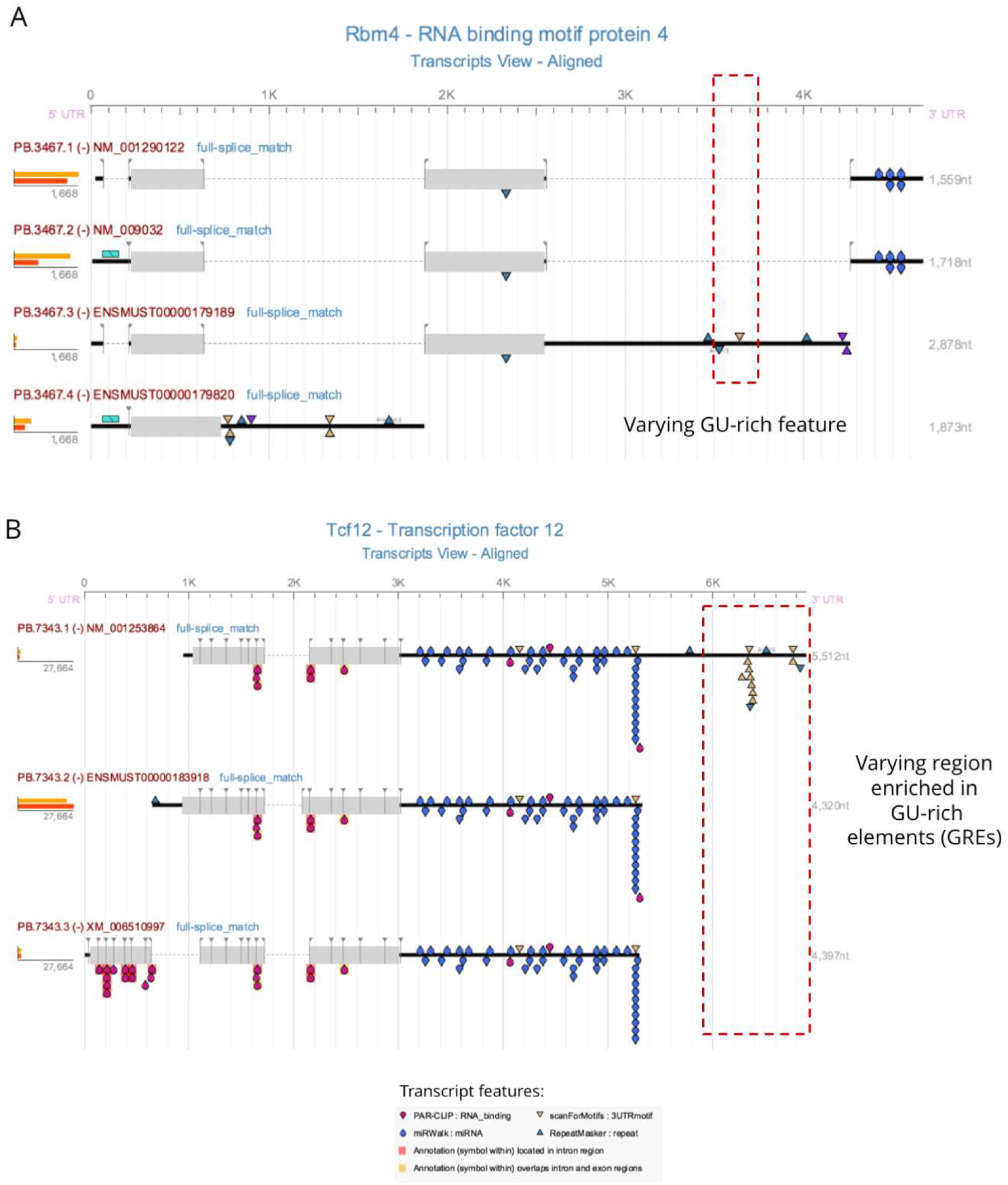
tappAS visualization of functional feature variation across isoforms. A) The *Rbm4* gene presents transcript-level variation in the inclusion of a GU-rich element (GRE) in the 3’UTR due to an exon-skipping event. B) Transcript-level variation in one of the isoforms of the *Tcf12* gene, which includes a 3’UTR region enriched in GREs due to an alternative Transcription Termination Site.

**Supplementary Figure 2:**
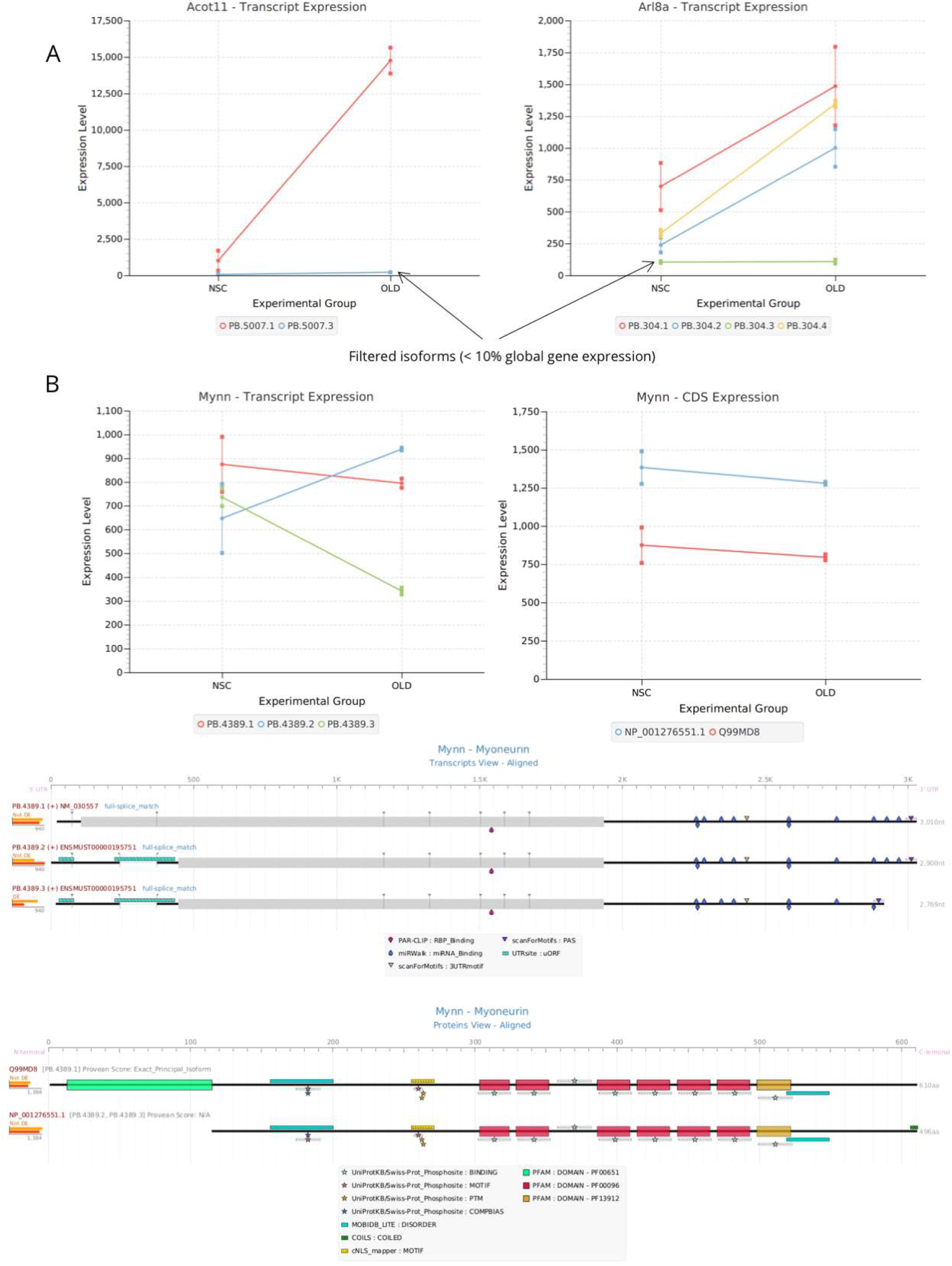
DE and DIU analysis results. A) Two examples of genes (*Acot11* and *Arl8*) detected as false positives for Differential Isoform Usage after minor isoform filtering (% expression < 0.1), i.e. where removal of the minor isoform leads to no DIU status. Filtered isoforms are indicated by arrows. B) Expression charts and tappAS visualization of annotated functional features at the transcript (left) and protein (right) levels for the *Mynn* gene, where Differential Isoform Usage and major isoform switching imply no Differential Coding sequence Usage.

**Supplementary Figure 3.**
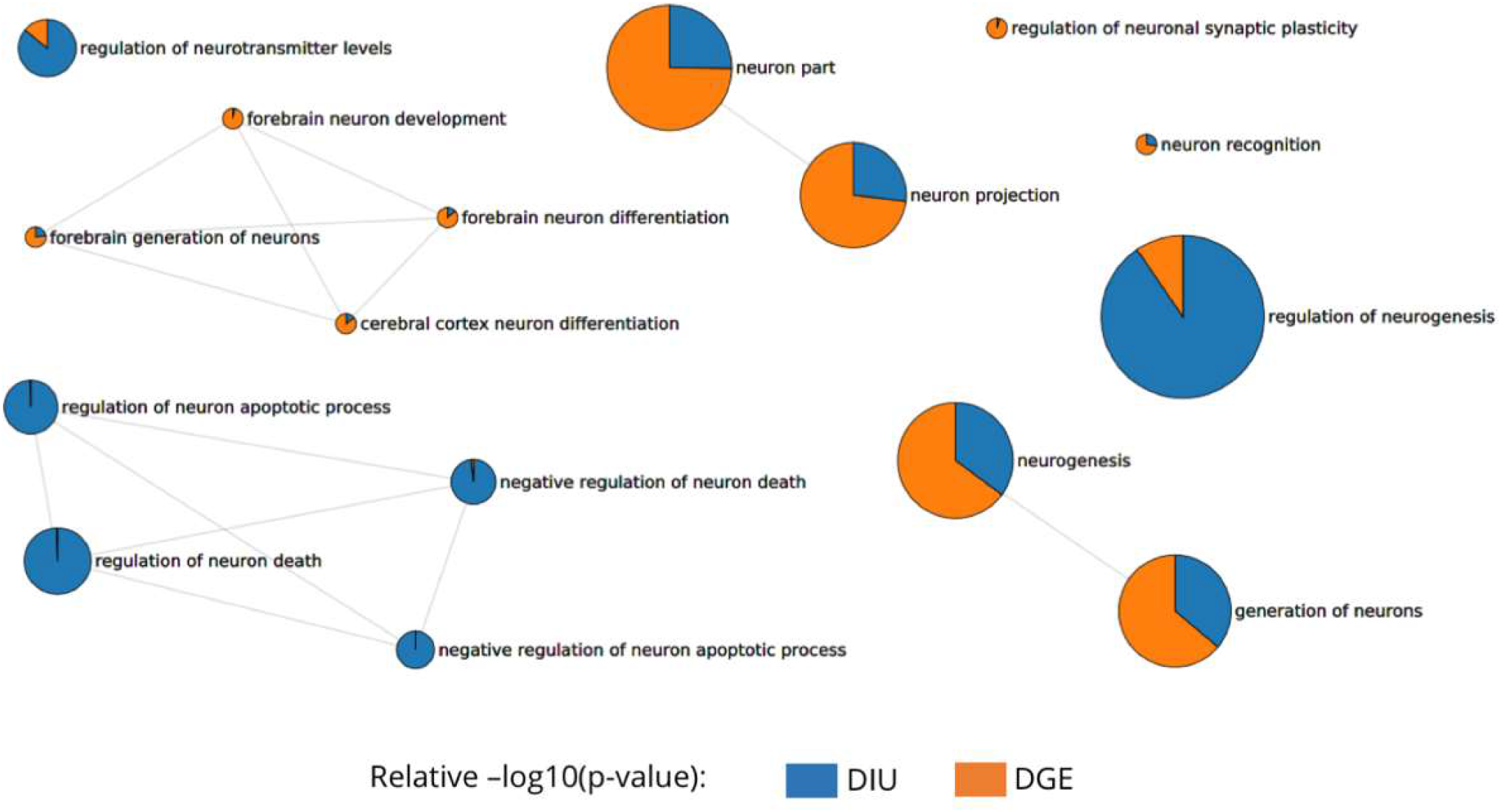
A) Clustering of GO term enrichment results for DE genes. Clusters are generated according to the similarities of significantly enriched GO terms. The color-coded legend indicates global process labels assigned after inspection of the different GO terms integrating each cluster. Important functions enriched in DE genes, i.e. affected by Differential Expression due to phenotypic differences between NSC and Oligodendrocytes, include ion/calcium homeostasis, cell motility and lipid metabolism. Circle size indicates enrichment Fisher Exact Test adjusted p-value. B) Relative functional relevance between DE and DIU regulation obtained in Multi-Dimensional Gene Set Enrichment Analysis of DE and DIU genes, representation of neural-related terms. Nodes correspond to GO-terms obtained by selecting the top-10 terms ranked by significance in the DE enrichment and the top-10 terms ranked by significance in the DIU enrichment (from a list of all neural-related GO-terms). Pie chart area represents DE and DIU regulation, and corresponds to relative −log 10(p-value).

**Supplementary Figure 4:**
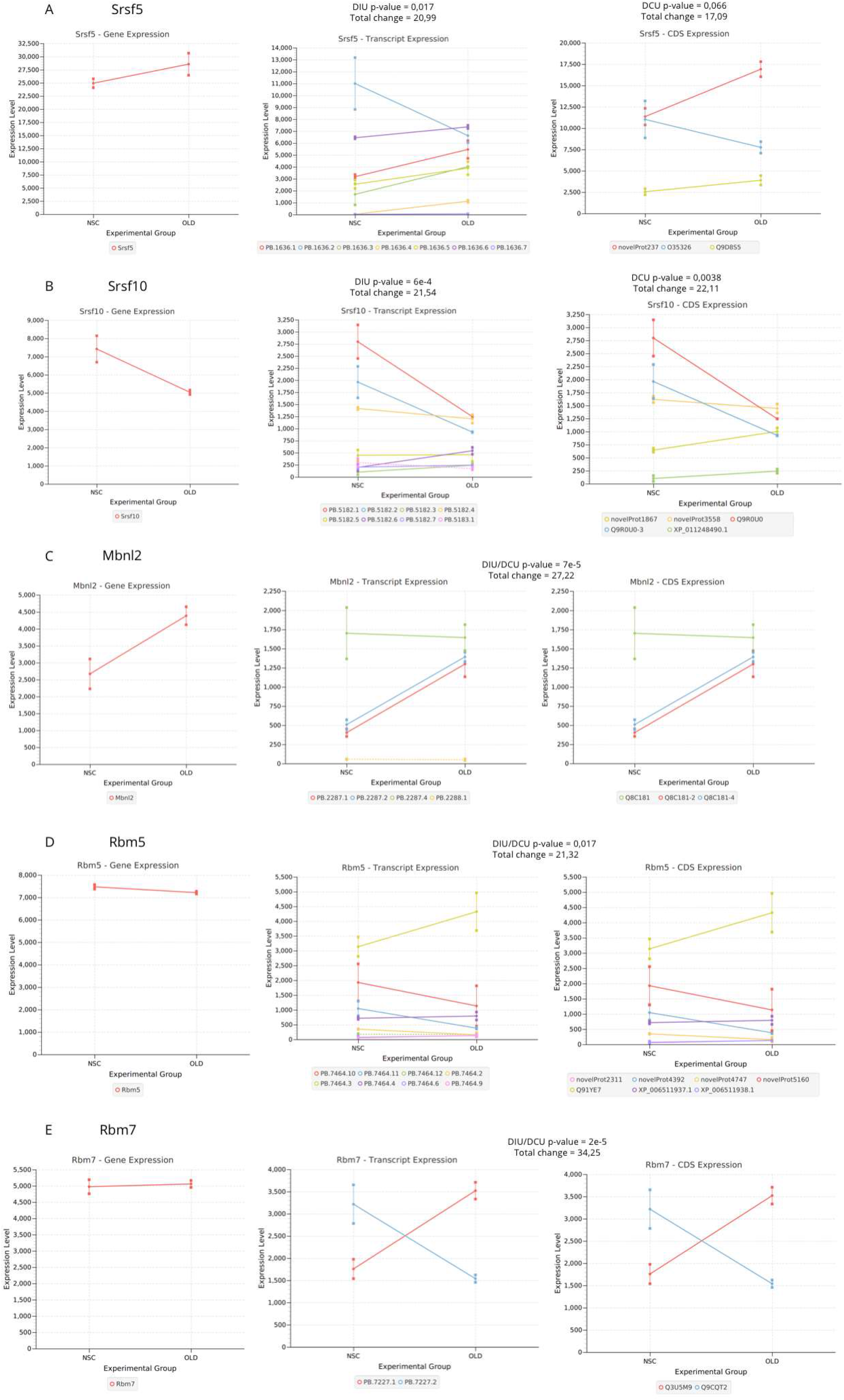
Splicing factors regulated by DIU. Transcript, gene and protein expression levels providing evidence of DIU status and self-regulation of the AltTP machinery: A) *Srsf5*, B) *Srsf10*, C) *Mbnl2*, D) *Rbm5*, E) *Rbm7*. DIU and/or DCU (indicated only when significantly different from DIU results) significance corresponds to multiple testing adjusted Q-Values.

**Supplementary Figure 5:**
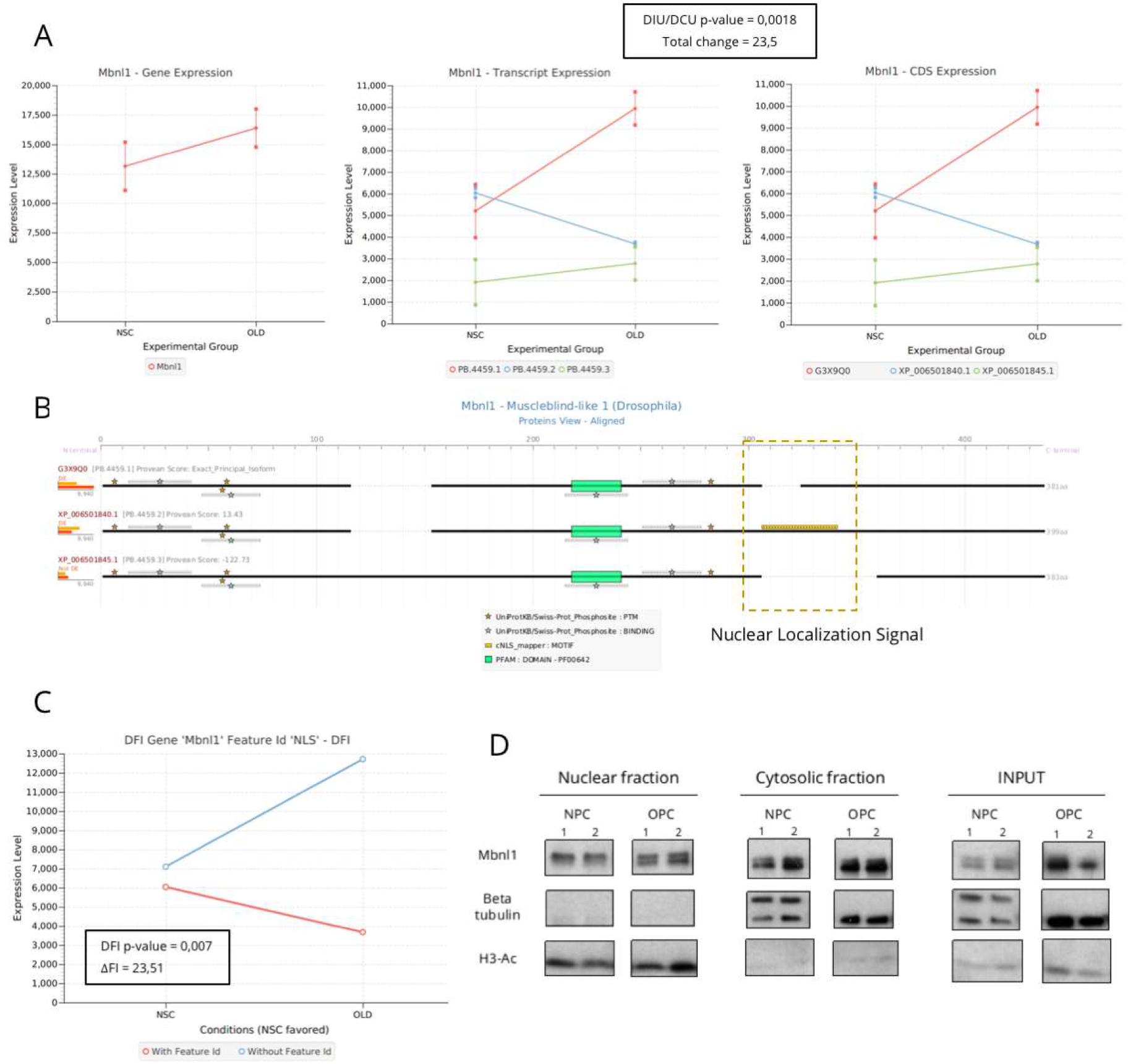
*Mbnl1* AltTP results. A) Gene, transcript and CDS expression for *Mbnl1*. The gene is positive for DIU both at the transcript and protein level. B) tappAS visualization of *Mbnl1* functional annotation. Differential inclusion of an NLS signal is detected by tappAS comprehensive annotation. C) DFI results for *Mbnl1* NLS signal. The feature is significantly differentially included, and favoured in NPCs. D) Western blot analysis of *Mbnl1* in cytosolic and nuclear fractions of NPCs and OPCs. Together with a general increase in *Mbnl1* expression in OPCs (INPUT), an increase in protein levels in the cytoplasm is observed, likely due to exclusion of the NLS signal (Cytosolic fraction).

**Supplementary Figure 6:**
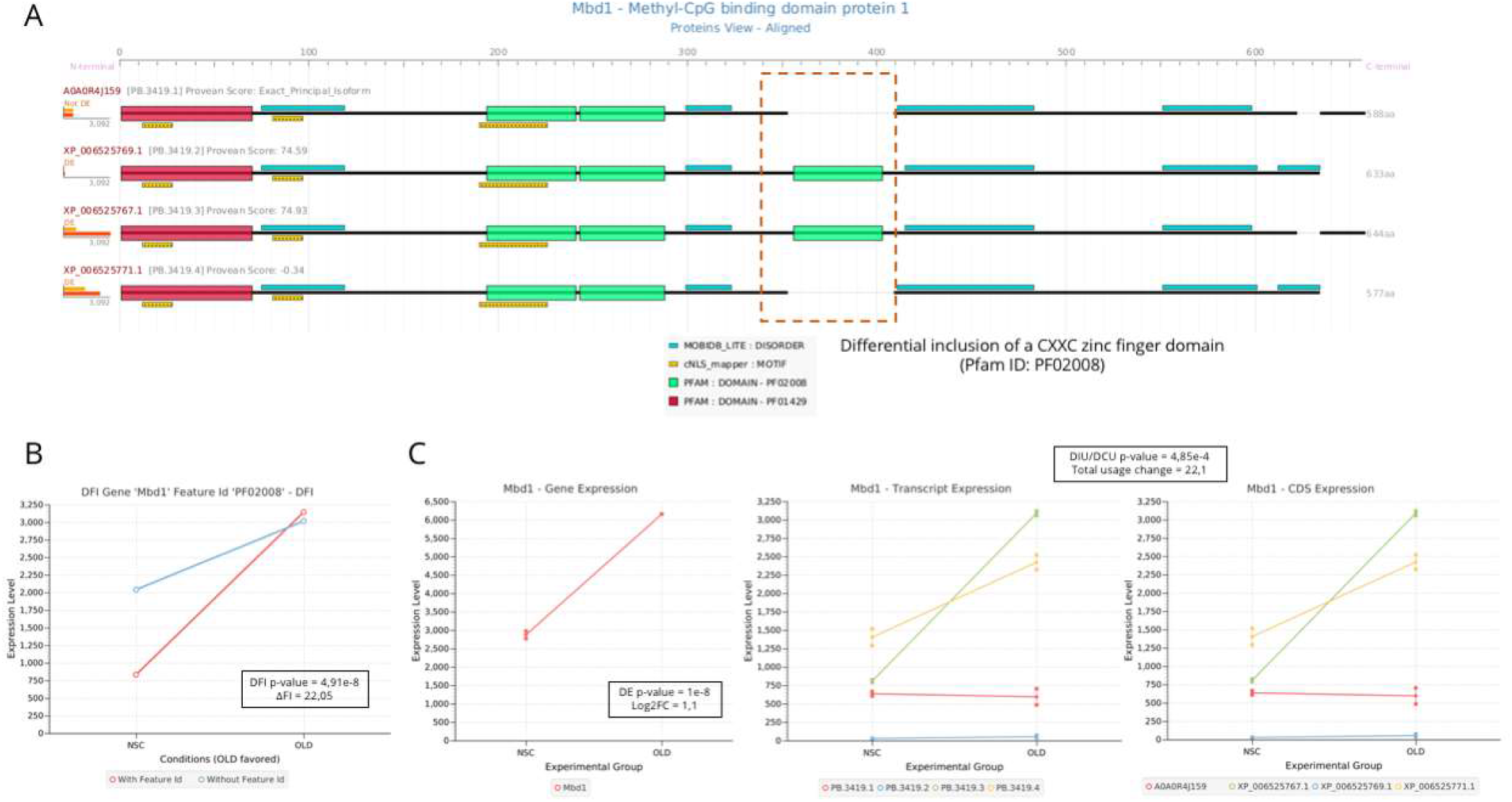
potential AltTP regulation of Mbd1 in DNA binding properties. A) Protein-level functional features annotated in tappAS. Highlighted area indicates differential inclusion of a third CXXC zinc finger domain to the coding region. B) DFI results for the CXXC zinc finger domain in the *Mbd1* gene. Inclusion of the domain, together with a general upregulation of *Mbd1*, are observed. C) Gene, transcript and CDS-level expression of *Mbd1*. The gene presents DE, DIU and DCU status.

**Supplementary Figure 7:**
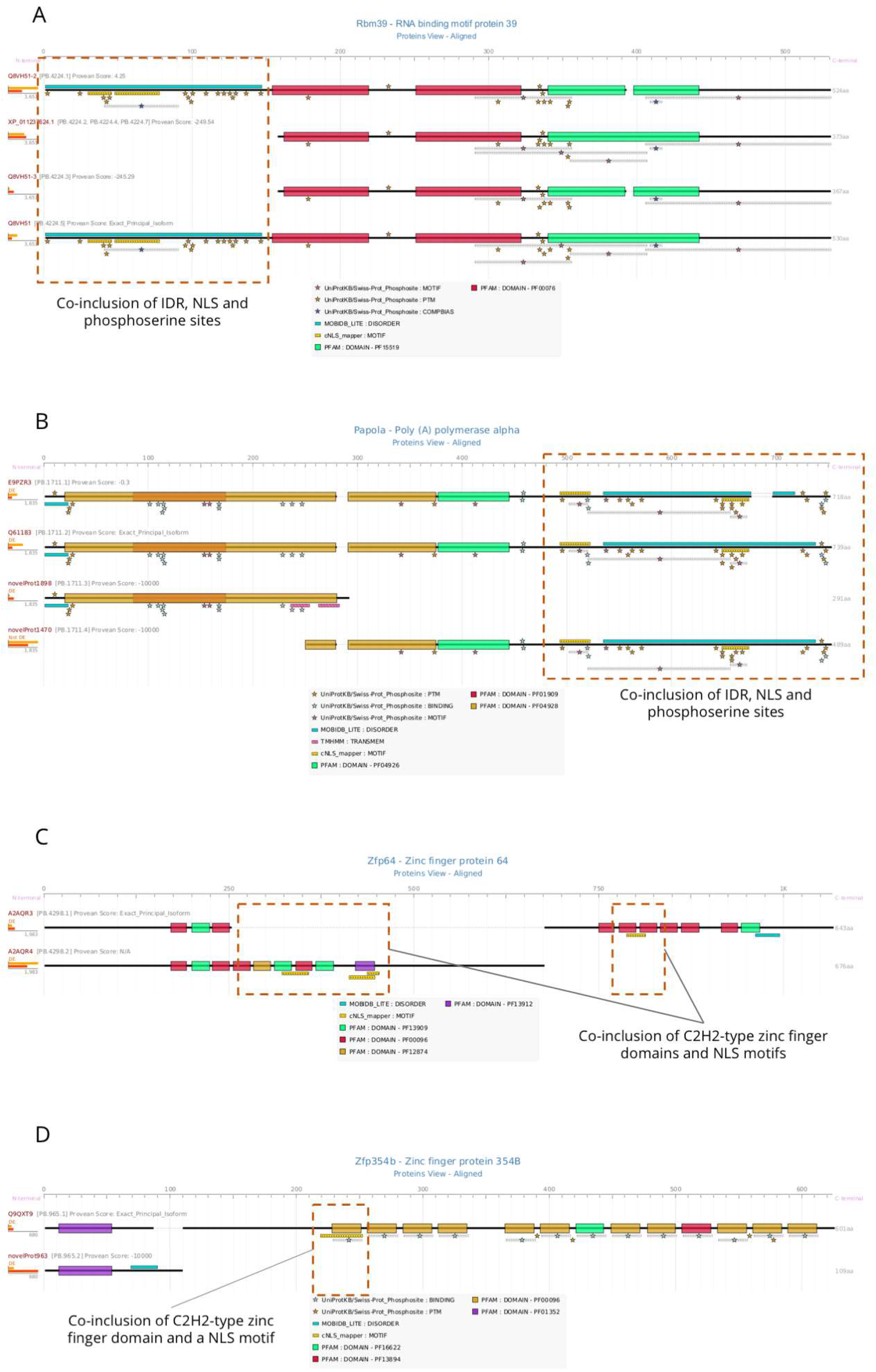
co-DFI results examples. A) Variation in the inclusion of protein-level functional elements in the *Rbm39* gene, which presents co-DFI status for an Intrinsically Disordered Region (IDR, DISORDER), several phosphoserine residues (PTM) and a Nuclear Localization Signal (NLS, MOTIF). B) Protein-level functional elements in the *Papola* gene, which presents co-DFI status for an IDR (DISORDER), several phosphoserine residues and two NLS (MOTIF). C) Protein visualization of he *Zfp64* gene, which presents co-DFI status of several C2H2-type zinc finger domains (PF13912, PF00096 and PF13909) and NLS motifs. D) The *Zpf354b* gene presents co-DFI status of a C2H2-type zinc finger domain (PF00096) and an NLS motif.

**Supplementary Table 1:** summary of annotation results for the mouse transcriptome of NPC and OPC primary cells. Number of features at the transcript and protein levels annotated are indicated, together with their database of origin and the percentage of isoforms in the transcriptome that contain them.

| Annotation level | Source | Category | No. feature occurrences | No. isoforms annotated |
| --- | --- | --- | --- | --- |
| Transcript<br>11970 isoforms<br>7167 genes | ScanForMotifs | PAS | 8511 | 5750 (48%) |
|  | ScanForMotifs | 3'UTR motifs | 11797 | 5325 (44%) |
|  | UTRscan/UTRsite | 5'UTR motifs | 325 | 315 (3%) |
|  | UTRscan/UTRsite | uORF | 7444 | 3045 (25%) |
|  | RepeatMasker | Repeat regions | 19269 | 7245 (61%) |
|  | MiRWalk/miRbase + in-house scripts | 3'UTR miRNA binding sites | 106392 | 9474 (79%) |
|  | clipDB + in-house scripts | RNA-binding sites (RBPs) | 47821 | 7279 (61%) |
| Protein<br>10813 coding isoforms<br>7167 genes | In-house scripts | Nonsense-Mediated Decay (NMD) | 329 | 329 (3%) |
|  | PFAM-HMMER3 | Domains | 20973 | 9608 (89%) |
|  | COILS + UniprotKB | Coiled coil | 6669 | 2856 (26%) |
|  | TMHMM + UniprotKB | Transmembrane regions | 12543 | 2061 (19%) |
|  | SignalP | Signal peptides | 824 | 824 (8%) |
|  | MOBIDB | Disordered regions | 11256 | 5626 (52%) |
|  | cNLS mapper + UniprotKB | Nuclear Localization Signals (NLS) | 7599 | 4297 (40%) |
|  | PSP + UniprotKB | Post-Translational Modifications (PTM) | 100804 | 8506 (79%) |
|  | UniprotKB | Compositional bias | 2260 | 1480 (14%) |
|  | UniprotKB | Motif | 6579 | 2897 (27%) |
|  | UniprotKB | Intramembrane | 159 | 62 (0.6%) |
|  | UniprotKB | Active site | 1770 | 1168 (11%) |
|  | UniprotKB | Binding | 12790 | 3339 (31%) |

**Supplementary table 2:** ID-level FDA results for UTR motifs, top 5 ranked by adjusted p-value. Significance assessed via Fisher’s Exact Test with Bonferroni-Hochberg multiple-testing correction.

| **3' UTR motif** | **p-value** | **Adj.p-value** | **No. varying genes (%)** |
| --- | --- | --- | --- |
| GU-rich Destabilization Element | 0.0024 | 0.05 | 197 (69.4%) |
| GU-Rich Element (GRE) | 0.0281 | 0.33 | 243 (66.2%) |
| tRNA-like structure | 0.079 | 0.53 | 279 (64.7%) |
| Brd-Box | 0,088 | 0.53 | 189 (65.4%) |
| Dinucleotide Repeat | 0.283 | 1 | 32 (66.6%) |

**Supplementary table 3:** ID-level FDA results for miRNA binding motifs, top-5 ranked by adjusted p-value. Significance assessed via Fisher’s Exact Test with Bonferroni-Hochberg multiple-testing correction.

| **miRNA** | **p-value** | **Adj.p-value** | **No. varying genes (%)** |
| --- | --- | --- | --- |
| mmu-miR-335-3p | 9e-4 | 0.46 | 61 (62.2%) |
| mmu-miR-590-3p | 0.0059 | 0.77 | 83 (56.8%) |
| mmu-miR-880-3p | 0.0071 | 0.77 | 21 (70%) |
| mmu-miR-7b-3p | 0.0101 | 0.77 | 43 (60.5%) |
| Mmu-miR-223-3p | 0.0145 | 0.77 | 35 (61.4%) |

## Methods

### Retrieving isoform-resolved functional annotation features

tappAS uses a gff3 like file with transcript structural and functional data. To produce this file for our mouse data (Supplementary Table 1) we use available databases and state-of-the-art prediction algorithms. Features are gathered through two mechanisms: positional transfer from functional databases and *de novo* prediction by state-of-the-art algorithms for sequence-based function prediction. All functional labels annotated at the isoform resolution are positionally described via their exact localization within protein/RNA molecules.

RNA-level annotations included: cis-acting UTR regulatory elements and Upstream Open Reading Frames (uORFs) predicted by UTRscan^87^; repeat regions and low-complexity elements predicted by repeatMasker^88^; and miRNA binding sites collected from mirWalk2.0^89^. A minimum seed length of 7bp and a p-value threshold of 0.05 were set as requirements to call miRNA binding sites. We filtered the site list by the number of sources reporting the association, requiring that miRNA binding sites to be predicted by a minimum of 5 methods, among which Targetscan^90^, miRanda^91^, and mirWalk^89^ are required. mirWalk provides transcript coordinate information to locate miRNA binding sites. High confidence miRNAs can be identified using the experimental evidence information in miRBase^92^. In our example there were 511 miRNAs with annotated binding sites and experimental evidence. Binding sites for RNA-binding proteins (RBPs) can be annotated by collecting genomic crosslinking immunoprecipitation (CLIP) data from CLIPdb^93^ and mapping sites to isoforms. RNA binding sites can be transferred by user defined levels of stringency. For our example, we required prediction by at least two algorithms in CLIPdb.

At the protein level, Pfam domains are mapped with InterProScan^94^, transmembrane regions predicted with TMHMM^95^, signal peptides obtained by SignalP 4.0^96^, coiled-coil regions predicted by COILS^97^, single and bipartite Nuclar Localization signals mapped by cNLS mapper^98^ (score > 6) and disordered regions obtained by MobiDB Lite^99^, which derives consensus IDR predictions by combining 8 different predictors. We predicted isoforms containing a premature termination codon (PTC)-potentially leading to nonsense-mediated decay (NMD)-using the 50-NT rule^100^ that indicates that a termination codon situated more than 50-55 nt upstream of an exon-exon junction is generally a PTC.

In addition to sequence-based prediction methods, some protein-centric databases contain a detailed annotation of protein features. However, these are generally biased towards the annotation of the best-documented isoform, hindering the study of the functional diversity of alternative isoforms. To correct this, we map canonical isoform annotations to query isoform sequences, novel or known, following an isoform-aware positional transfer strategy. We obtained the information on protein functional features by parsing UniprotKB^101^ and PhosphoSitePlus^102^ databases. In both cases we deal with the disparities between databases when defining gene models and ensure the ORF and genomic position conservation between public and query sequences during feature transference. As a result, we retrieved an extensive set of post-translational modification (PTM) sites with experimental evidence from PhosphoSitePlus, and a diverse catalogue of functional sequence features from UniprotKB.

tappAS contains precomputed gff3 files with isoform functional data for mouse, human, Arabidopsis, fly and maize. Specific details can be found in Supplementary Table 1.

### Visualization engine of positional functional annotation at isoform resolution

The tappAS visualization engine is designed to display isoform variability in a user-friendly manner, and constitutes one of the most useful features of the application. Using the visualization power of the Java engine, tappAS displays the whole catalogue of isoform-resolved annotation features and their position using a distinctive icon on both transcript and protein isoform structure maps. Maps include UTR/CDS areas, polyA sites, splice junction and exon information, and functional features, creating a graphical representation that greatly facilitates the study and comparison of isoform diversity.

### Functional Diversity (FD) Analysis

Isoforms vary in structural and functional features among isoforms of the same gene. FD identifies and measures the nature of the variability in a qualitative manner. For every annotated feature, all pairwise comparisons between transcript isoforms from the same gene are performed and a gene is labelled as *varying* if at least one isoform pair has variability in a feature, either in its annotated genomic position(s) (*Positional Varying*) or in the presence/absence of the annotated feature (*Presence Varying*). Functional Diversity can be assessed by gene or by feature ID.

#### Gene-Level Diversity

The Gene-Level Diversity analysis evaluates genes as a function of the structural, functional and regulatory feature categories that are modulated by AltTP. Depending on the feature category and its relationship to the functional properties of a transcript or protein, Functional Diversity is evaluated using a *Positional Varying* strategy or a *Presence Varying* strategy.

The *Positional Varying* approach compares features by genomic position, i.e. by mapping features to genomic coordinates and classifying them as varying if coordinates are not equivalent between gene isoforms. Position disagreement is annotated when >9bp, that is, 3 amino acids, allowing for variability in prediction. In contrast, *Presence Varying* includes only presence/absence of annotation. For instance, NMD transcript status is is, based differences in the transcript level NMD label. In contrast, transcript attributes such as UTR length, CDS and polyA site positions, are examples of features where *positional* evaluation is meaningful. However, a third group of features (such as Pfam domains or transmembrane regions) can be affected by AltTP via both complete and partial disruption of the feature. In these and similar cases, both strategies can be used, and provide complementary insight on AltTP in the potential regulation of the functional or regulatory feature.

For structural features evaluated by *Positional Varying*, some special considerations are required. In order to detect alternative polyadenylation (APA) events, polyA sites are identified as the last genomic position of transcript isoforms and evaluated in a pairwise manner by computing the polyA distance between each pairwise combination of isoforms expressed by a given gene. mRNA cleavage is not an exact process and can occur within a small window of positions^103^. To take cleavage variability into account, tappAS’ FD analysis labels a pair of isoforms as APA when there is a minimum X bp genomic distance (default value 100) between polyA sites. UTR length is computed for each isoform for subsequent pairwise comparison between coding isoforms from the same gene. Pairs of isoforms with 3’/5’ UTR differences above a user-specified cutoff (75 bp by default) are labelled as 3’/5’ UTR length varying, respectively. Finally, CDS variability is determined by comparing CDSs both at the sequence and genomic coordinate levels. Non-coding isoforms are discarded from CDS diversity analysis.

#### Feature-Level Diversity

The Feature-Level Diversity analysis identifies specific functional and regulatory elements (i.e. by feature ID instead of source/functional category) varying across isoforms from the same gene. Feature-Level Diversity Analysis counts the number of genes for which a given feature ID is flagged as *varying* in the gene level analysis. The diversity status of each ID can be evaluated via *Positional, Presence Varying* or both.

The significance level of every feature global variation across genes is evaluated using Fisher’s Exact Test^104^, and then corrected using the Benjamini-Hochberg^105^ method for multiple testing correction.

### Differential Feature Inclusion (DFI) analysis

DFI applies the concept of exon inclusion analysis to functional features. DFI is only applied to features labelled as varying –either by position or as present/absent-across each gene’s isoforms, as only these have the potential to be significantly regulated. For a given gene and functional element, the null hypothesis that transcripts containing the feature have equivalent expression to transcripts not containing the feature is tested for each gene. Expression values of the isoforms containing the feature, and isoforms where the feature is not present are calculated from the data.

The *feature inclusion rate* is the ratio between the sum of expression of all feature-including isoforms and the total expression of the gene (i.e. sum of expression of isoforms including and excluding the feature) for each condition studied:

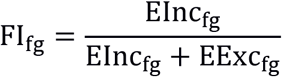

where *EInc* is the aggregated expression value for feature-including isoforms and *EExc* is the aggregated expression value for feature-excluding isoforms for gene *g* and positional feature *f*.

Differential inclusion of functional features is then tested using generalized linear models adapting DEXSeq^6^ and maSigPro^106^ methods, for case-control and time-course experimental designs, respectively. For each feature *f* and gene *g*:

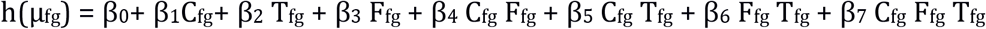

where *h* is the link function of the GLM, μ_fg_=E(y_fg_) is the expected aggregated expression level, *C_fg_* is the binary variable that identifies each of the two experimental conditions, T_fg_ is the time point, and *F_fg_* is the binary variable that identifies the variant (Feature-Excluding or Feature-Including).

Each gene and feature are individually modeled. For each model, the significance of the condition-variant or condition-variant-time interactions is evaluated, depending on the experimental design considered. When multiple functional annotation categories are analyzed (domains, UTR motifs, disordered regions, etc.), each of them is tested independently. P-values are corrected by FDR and significance is set to 0.05 by default.

### Co differential feature inclusion analysis (Co-DFI)

Co-DFI analysis evaluates how frequently two features are simultaneously DFI for the same gene in the same condition, while mutual exclusion evaluates how often two features are simultaneously DFI for the same gene in the different condition. Co-DFI is computed for each pair of features detected as DFI in at least 5 genes.

### Defining a library of polyA sites

tappAS uses a polyA site database is created by extracting the genomic coordinate of the last position of each transcript isoform. Unlike recently developed tools^107^, polyA sites in terminal exons with different 5’ start sites are also considered to allow the analysis of Alternative PolyAdenylation sites affecting either Coding (CR-APAs) or UTR- (UTR-APAs) events. Non-coding isoforms as well as NMD-predicted variants are discarded.

Next, a series of filtering and collapsing steps are performed in order to define the proximal (pPA) and distal polyA (dPA) site for each gene. First, independent cleavage sites are defined by merging polyA sites located within a 75 bp window. To avoid the definition of a minor polyA site as a distal or proximal site, a filtered based on relative polyA site expression levels is applied and only polyA sites accumulating at least 10% (default threshold) of total gene expression in at least one condition are considered. In the case of genes with more than two polyA sites, we perform a final merge of unlabelled sites by assigning them to the nearest proximal or distal site.

### Differential Polyadenylation Analysis (DPA)

Using the defined polyA site library, tappAS computes the per-gene and per-sample dPA and pPA site expression levels by collapsing the expression levels of the set of transcript isoforms that contain either the dPA or de pPA. The same GLM model used for DFI is applied to capture significant condition-variant interactions. The relative distal polyA site usage (DPAU) is implemented by calculating the relative expression of the sum of all isoforms containing the distal site over the total polyA site expression level of the gene:

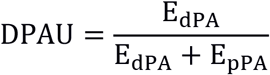

where *EdPA* and *EpPA* to the expression levels of the variants defined as distal and proximal polyA sites.

### Detecting lengthening and shortening of 3’ UTRs

For isoforms with identical CDS end positions but different polyA (UTR-APAs) distal/proximal polyA site usage directly associates with UTR lengthening/shortening events. However, when changes in polyA site position imply changes in the CDS (CR-APAs), it is impossible to directly infer the relationship between the polyA site and 3’ UTR length. Since DPA analysis assesses polyA site regulation independently of the coding sequence, tappAS introduces a specific 3’ UTR lengthening/shortening analysis by computing an isoform usage-weighted mean UTR length for each condition:

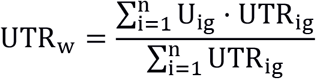

where *U* is the relative usage of isoform i in gene *g* and *UTR* its associated 3’ UTR length.

UTRs from highly expressed isoforms will contribute in a higher proportion to the final UTR mean length. The weighted UTRs is a measure of the actual extent of UTR length changes across conditions. Statistical differences are tested by using a Wilcoxon rank-sum test of the weighed UTR values.

### tappAS software

tappAS (http://tappas.org) is a Java GUI application that provides a broad analytical framework including a range of functions that, collectively and in combination allow the study of different structural and functional aspects associated to isoform usage. Statistical methods are implemented in R and are run by tappAS using the Rscript environment. See http://tappas.org for a comprehensive list of R package dependencies, as well as other software and hardware requirements.

tappAS uses a gff3-like file containing functional annotations at the isoform level, as described above. Structural information, including annotation of UTRs, CDS and introns, together with gene, protein and transcript reference IDs for each transcript sequence are also represented in the this gff3-like file. Currently, annotation files for Human (ENSEMBL and RefSeq databases), Mouse (ENSEMBL and RefSeq databases), Drosophila, Arabidopsis and Maize are available in the tappAS application. Users can optionally input their gff3 files.

tappAS works as a compendium of independent projects, each of them created using two inputs: a transcript expression matrix and an experimental design file, that can either be a case-control or a time-course experiment. Being a GUI application, tappAS also provides a rich set of interactive features via the JavaFX platform, including customizable data tables, complex sorting and filtering options, data and figure export, context-sensitive help pages, data drill-down and display customization.

tappAS is designed an open framework and accepts user-defined gene lists as input for analysis.

While tappAS accepts a transcript-expression file, it uses the structural annotation included in the gff3 to estimate the expression of genes, as the sum of their transcript expression, and CDSs, as the sum of transcripts having the same ORF. Differential expression analyses can be then run at each of these aggregation levels.

The application includes the analysis methods described above and implements existing tools when appropriate, including NOISeq^108^ and maSigPro^106^ for differential gene expression; DEXseq^6^ and Iso-maSigPro^109^ for differential isoform usage. These two methods assess DIU by fitting generalized linear models (GLMs) and testing the significance of the isoform-condition interaction coefficient, as proposed in^110^. Implemented enrichment methods are GOSeq^58^ (Functional Enrichment) GOglm^59^ (Gene Set Enrichment) and mdgsa^69^ for multi-dimensional GSE. These enrichment tools can be easily applied to the results of any of the statistical methods included in tappAS. Finally, tappAS implements extant complementary functionalities (i.e. low-count expression filtering, TMM normalization, PCA, clustering methods) that enable the pre-processing and flexible exploration of data and results.

### Complementary metrics for isoform analysis

tappAS incorporates complementary analysis features and metrics specially conceived for a better assessment of the functional implications of AltTP.

#### a) Major and minor isoforms

In tappAS, the major isoform of a gene is defined as the isoform with the highest mean expression across all conditions of the study, while other isoforms of the gene are labelled minor forms. Such definition is operational in the context of tappAS analysis and does not assume functional relevance or expression levels in other experimental settings.

#### b) Isoform prefiltering

Genes in mammalian transcriptomes usually express multiple of isoforms. However, frequently only one or few of them accumulate the major proportion of gene expression^111^ while remaining isoforms, although detected, have low expression levels. Although tappAS allows for low expression filtering upon data upload, still isoforms may remain that are relatively minor for their gene expression level. When the minor isoforms have small expression changes between conditions, but these occur in the opposite direction to the predominant isoforms, significant isoform-condition coefficients may appear at GLM models. To avoid the detection of DIU genes because of the ‘flat’ behaviour of minor isoforms, an isoform filtering step can be applied before statistical modeling. Two filtering approaches are implemented in tappAS. One considers the proportion of a gene’s expression represented by each isoform and filters those that do not reach a minimum expression rate (10% by default), while the other calculates the fold-change of the minor isoforms versus the major to remove those below a specified fold-change (FC) threshold (default FC=2). Users can use which filtering option to apply

#### c) Total usage change

##### Total usage change in DIU analysis

The Fold Change is a measure magnitude in of differential expression that cannot be easily applied to DIU analysis, where multiples isoforms are tested in a single model. Instead, we propose a new metric, *total usage change* to quantify the magnitude of change for DIU genes. *Total usage change* measures the amount of redistribution (as %) in expression levels across different conditions for isoforms of the same gene. Because absolute gene expression levels may be different across conditions, total change values are always represented as a function of the gene expression FC.

We define isoform usage as the relative expression of isoform *i* in gene *g*. Then, total usage change can be defined as:

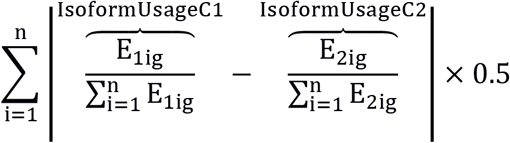

where *E_ig_* is the expression value for isoform *i* and gene *g*.

##### Defining total usage change for feature analyses

When performing DFI and DPA analyses, expression values are collapsed to obtain feature inclusion (FI) and distal polyadenylation site usage (DPAU) levels. In this case, total change is re-defined as the redistribution (as %) of FI (ΔFI) or DPAU (ΔDPAU) levels across every pair of conditions considered. Note that both metrics are also dependent on absolute expression.

#### d) Defining switching events

*Switching events* are defined in order to identify Differential Feature Inclusion, Differential Polyadenylation and Differential Isoform Usage events that imply a strong change and prioritize candidates for further analysis.

A *major isoform switching event* occurs when the major isoform of gene the becomes minor at one particular condition. In multiple time-course series major isoforms are defined for each experimental group and so is the major isoform switch. *Feature switching* (in DFI) and *distal polyA usage switching* (in DPA) are similarly defined.

##### Favored conditions

In DPA and DFI analyses, switching information is complemented by information of the favored condition, i.e. the experimental condition where the inclusion of the feature is promoted.

### Experimental setup in murine neural cells

As proof-of-concept of our analysis framework, we used the data in Tardaguila et al^57^. Briefly, two different cell types: Neural Precursor Cells (NPC) and Oligodendrocyte Progenitor Cells (OPCs) had expressed transcriptomes estimated using PacBio Iso-Seq sequencing and curated by SQANTI^57^, resulting in 11,970 transcripts coded by 7,167 genes. We computed isoform expression levels with RNA-seq using RSEM^112^ following ENCODE guidelines

### Validation of events with potential functional impact

#### Western Blot

We validated AltTP-mediated localization changes via differential inclusion of Nuclear Localization Signals (Ctnnd1, Mbnl1) using nucleus-cytoplasm fractioning and western blot analyses. Total protein fraction was extracted from cell cultures using lysis buffer containing 50 mM Tris-HCl, pH 7.5,150 mM NaCl, 0.02% NaN3, 0.1 SDS, 1% NP40, 1 mM EDTA, 2 mg/mL leupeptin, 2 mg/mL aprotinin, 1 mM PMSF, 1x Protease Inhibitor Cocktail (Roche Diagnostics, San Diego, CA, USA). The cytoplasmic and nuclear protein fractions were extracted with lysis buffer containing HEPES 10 mM pH 7.9, KCl 10 mM, EDTA 1 mM, EGTA 1 mM, DTT 1 mM, B-glycerophospate 10 mM and 1x Protease Inhibitor Cocktail (Roche Diagnostics, San Diego, CA, USA). IGEPAL (CA-630) 0.4% was then added and samples were vigorously vortexed and centrifuged at 12000g at 4°C for 5 minutes. The supernatant (cytoplasmic fraction) was recovered, and the remaining pellets were incubated in lysis buffer containing 10 mM TRIS pH 7.4, NaCl 400 mM, IGEPAL (CA-630) 0.5%, EDTA 1 mM, EGTA 1 mM,DTT 1 mM, B-glycerophospate 10 mM and 1x Protease Inhibitor Cocktail (Roche Diagnostics, San Diego, CA, USA) to recover nuclear protein extracts (nuclear fraction). The protein concentrations of the supernatant were determined via bicinchoninic acid technique (Pierce^®^ BCA protein assay; Thermo Fisher Scientific) and stored at −80C. Equal protein amounts were loaded, separated in 10% SDS–PAGE and transferred into a PVDF membrane. The membrane was blocked with 5% milk in TBS with 0.1% Tween-20 for 1h at room temperature and incubated at 4 °C overnight with the following primary antibody solutions (4% milk, 0,5% Tween-TBS): Anti p120 (Ctnnd1) 1:2000 (Millipore 05-1567, clone 15D2); Anti Mbnl1 1:100 (DSHB-MB2a(3b4)); Anti Ac H3 1:1000 (Millipore 06-599); Anti tubulin coupled HRP (Thermo MA5-16308-HRP). Membranes were incubated for 1h at room temperature with the following secondary antibody dilutions (4% milk, 0,5% Tween-TBS): Anti mouse HRP (Life A16072) 1:10000; Anti rabbit HRP (Thermo 31460) 1:10000. Signal detection was performed with an enhanced chemiluminescence kit (ECL Plus Western blotting detection reagent from GE Healthcare, Piscataway Township, NJ, USA) and bands were detected using Alliance Q9 Advanced (Uvitec Cambridge Inc).

